# Unraveling the relationship between cancer and life history traits in vertebrates

**DOI:** 10.1101/2022.07.12.499088

**Authors:** Stephanie E. Bulls, Laura Platner, Wania Ayub, Nickolas Moreno, Jean-Pierre Arditi, Saskia Dreyer, Stephanie McCain, Philipp Wagner, Silvia Burgstaller, Leyla R. Davis, Linda GR. Bruins - van Sonsbeek, Dominik Fischer, Vincent J. Lynch, Julien Claude, Scott Glaberman, Ylenia Chiari

**Affiliations:** Department of Biology, George Mason University, Fairfax, VA, USA; Department of Environmental Science & Policy, George Mason University, Fairfax, VA, USA; Der Grüner Zoo, Wuppertal, Germany; Allwetterzoo, Münster, Germany; Department of Biology, University of South Alabama, Mobile, AL, USA; Birmingham Zoo, Birmingham, AL, USA; Zoo Zurich, Zürich, Switzerland; Diergaarde Blijdorp, Rotterdam Zoo, Rotterdam, The Netherlands; Wildlife Zoo, Hellental, Germany; Department of Biological Sciences, University at Buffalo, SUNY, USA; Institut des Sciences de l’Evolution, Université de Montpellier/CNRS/IRD, Montpellier, France; Faculty of Science, Chulalongkorn University, Bangkok, Thailand; Centre for Environmental Research and Justice, School of Biosciences, University of Birmingham, UK; School of Life Sciences, University of Nottingham, Nottingham, UK

**Keywords:** Body mass, cancer prevalence, comparative oncology, intrinsic cancer risk, lifespan, neoplasia, phylogenetic comparative method, Peto’s paradox, vertebrates, zoo

## Abstract

Identifying species with unusually low cancer prevalence can provide new insights into cancer resistance. Most studies have focused on mammals, but the genetic, physiological, and ecological diversity among vertebrates can influence cancer susceptibility. We used necropsies from over a thousand species of amphibians, birds, crocodilians, mammals, squamates, and turtles to investigate relationships between cancer prevalence, intrinsic cancer risk, body mass, and lifespan. Previous studies often relied on species averages, leading to inaccurate interpretations. Our innovative statistical approach uses raw cancer data and resampling to improve accuracy. We found remarkably low cancer prevalence in turtles, high prevalence in squamates and mammals, and lower-than-expected prevalence based on lifespan and body mass in multiple groups. Our results show lifespan influences neoplasia and malignancy transformation rates in mammals, while body mass affects neoplasia prevalence in amphibians and squamates. These data reveal a complex relationship between life history traits and cancer risk, identifying vertebrates with potential novel cancer resistance mechanisms.

**STATEMENT OF SIGNIFICANCE:** Biodiversity is an untapped natural resource for understanding cancer. Our study reveals a wide divergence in cancer prevalence among vertebrate groups, with notably low rates in turtles and high rates in squamates and mammals. These findings can lead to new breakthroughs in understanding the biology of cancer.

## INTRODUCTION

Understanding cancer prevalence across animal species can identify novel cancer resistance mechanisms. Theoretically, larger, longer-lived species should have higher cancer susceptibility due to increased cell numbers and more extensive lifetime cellular turnover ^1^. However, empirical studies have not observed this pattern in mammals ^2–4^ or other vertebrates ^5,6^. This suggests that species with extreme body sizes and lifespans must have evolved mechanisms to offset their elevated cancer risk ^1,7,8^. Our study seeks to bridge this gap by investigating cancer prevalence within and among vertebrate groups, including underrepresented ones such as reptiles and amphibians, to identify species with novel anti-cancer mechanisms ^9^.

Prior research has documented marked differences in cancer prevalence among vertebrates, with higher rates in mammals compared to other groups ^10–14^. But studies have largely overlooked diversity within certain groups, especially reptiles and amphibians. There haven’t been any phylogenetic comparative studies of cancer that discriminate among major lineages of non-avian reptiles including crocodilians, squamates (lizards and snakes), and turtles. Previous data on cancer occurrence hint at low cancer prevalence in turtles and possibly crocodilians ^15,16^, underscoring the need for more in-depth analysis in these groups.

In this study, we assess cancer prevalence across all major vertebrate groups except fish—including amphibians, birds, crocodilians, mammals, squamates, and turtles—using zoo necropsy data. Our goal is to uncover the relationship between cancer prevalence and species-specific traits such as body mass, lifespan, and intrinsic cancer risk. We introduce an innovative statistical approach to overcome the significant limitations of previous research, employing a phylogenetic comparative method specifically designed for binomial distributions ^17^. This is crucial because the rarity of cancer within species results in prevalence data that are heavily skewed towards zero, fundamentally violating normality assumptions that many statistical models, including phylogenetic generalized least squares (PGLS) regression, depend on. Indeed, PGLS regression—commonly used in past studies ^3,6^, (see ^10^ for analysis of data using modified PGLS and the approach presented here) —relies on species averages, making it less appropriate for handling non-normal data characteristic of cancer prevalence.

In contrast, our innovative method leverages individual data points, offering a more precise account of intraspecific sampling effort and avoiding the common issue of treating prevalence as a continuous variable. This strategy establishes stronger correlations between cancer incidence and key biological traits across various vertebrate groups. Additionally, we incorporate a jackknife resampling technique to evaluate the robustness of our findings, particularly the impact of species sampling effort. This approach not only addresses the shortcomings of previous methodologies, but also provides a more reliable foundation for future research into cancer prevalence across vertebrates. Finally, we contextualize our findings within existing research by reanalyzing publicly available mammalian data using our improved statistical methodology ^3,4^ (see also ^10^ for a similar approach in comparing results obtained following the innovative approach used in this paper and a modified PGLS method).

Our analysis shows that neoplasia and cancer prevalence exhibit significant variation within and among vertebrate groups. Unlike previous studies ^2–6^, our results indicate that body mass and lifespan are influential factors for cancer prevalence in amphibians, squamates, and mammals. We further show that cancer prevalence is extremely low in turtles. This highlights the importance of employing precise statistical methods to reveal complex biological relationships and provides new perspectives on cancer biology across different vertebrate lineages.

## RESULTS

### Necropsy Data

Our dataset included 9,631 necropsy reports from 1,030 species (1,060 species, including additional squamate data from ^18^). Out of the total described species in each vertebrate group, our sampling represents 0.7% of all amphibians, 4.5% of all birds, 34.7% of all crocodilians, 4.0% of all mammals, 1.8% of all squamates, and 21.0% of all turtles (**Table 1**). Birds were highly represented with 507 species and 5,121 necropsy reports, while turtles and crocodilians had fewer necropsies (207 and 26, respectively). The Shannon Equitability Index indicated that the allocation of necropsies among species was similar within each group (**Supplementary Materials Table 2**).

**Table 1.** Total number of necropsies and species for each tetrapod group. The total recognized number of species for each group was obtained from IUCN (*The IUCN Red List of Threatened Species*, ^19^). “Total neoplasia” includes all benign and malignant tumor counts whereas “Total cancer” includes only those tumors that were diagnosed as malignant by a veterinary pathologist and confirmed by histology. Malignancy transformation rates is indicated as “malignancy prevalence” and was calculated as neoplasia count divided by cancer count.

|  | Amphibians | Birds | Crocodilians | Mammals | Squamates | Squamates* | Turtles |
| --- | --- | --- | --- | --- | --- | --- | --- |
| **Species in dataset** | 49 | 507 | 8 | 230 | 179 | 209 | 57 |
| **Recognized species in group** | 7296 | 11162 | 23 | 5968 | 9855 | 9855 | 269 |
| **Percentage of recognized species represented** | 0.7 | 4.5 | 34.7 | 4.0 | 1.8 | 2.1 | 21.0 |
| **Total necropsies** | 383 | 5121 | 26 | 2841 | 1053 | 2608 | 207 |
| **Total neoplasia** | 15 | 96 | 2 | 191 | 84 | 198 | 1 |
| **Total cancer** | 10 | 61 | 1 | 111 | 57 | 164 | 0 |
| **Neoplasia prevalence** | 0.04 | 0.02 | 0.08 | 0.07 | 0.08 | 0.08 | 0.005 |
| **Cancer prevalence** | 0.03 | 0.01 | 0.04 | 0.04 | 0.05 | 0.06 | 0 |
| **Malignancy prevalence** | 0.7 | 0.6 | 0.5 | 0.6 | 0.7 | 0.8 | 0 |

\*Includes data from <sup>18</sup>

### Prevalence of Neoplasia, Cancer, and Malignancy Transformation Rate

We found clear differences among tetrapod groups (**Figures 1**, **2**, and **3**), with some groups having notably low neoplasia, cancer prevalence, and/or malignancy transformation rates. Turtles had no instances of cancer and only one case of neoplasia. Crocodilians had only two neoplasms, one of which was malignant (**Table 1**). Neoplasia and cancer prevalence values were also particularly low in birds (2% and 1%, respectively), while the malignancy transformation rate was 60% (**Table 1**). These results indicate that while cancer is relatively rare in birds, it is frequently malignant when it does occur.

**Figure 1.**
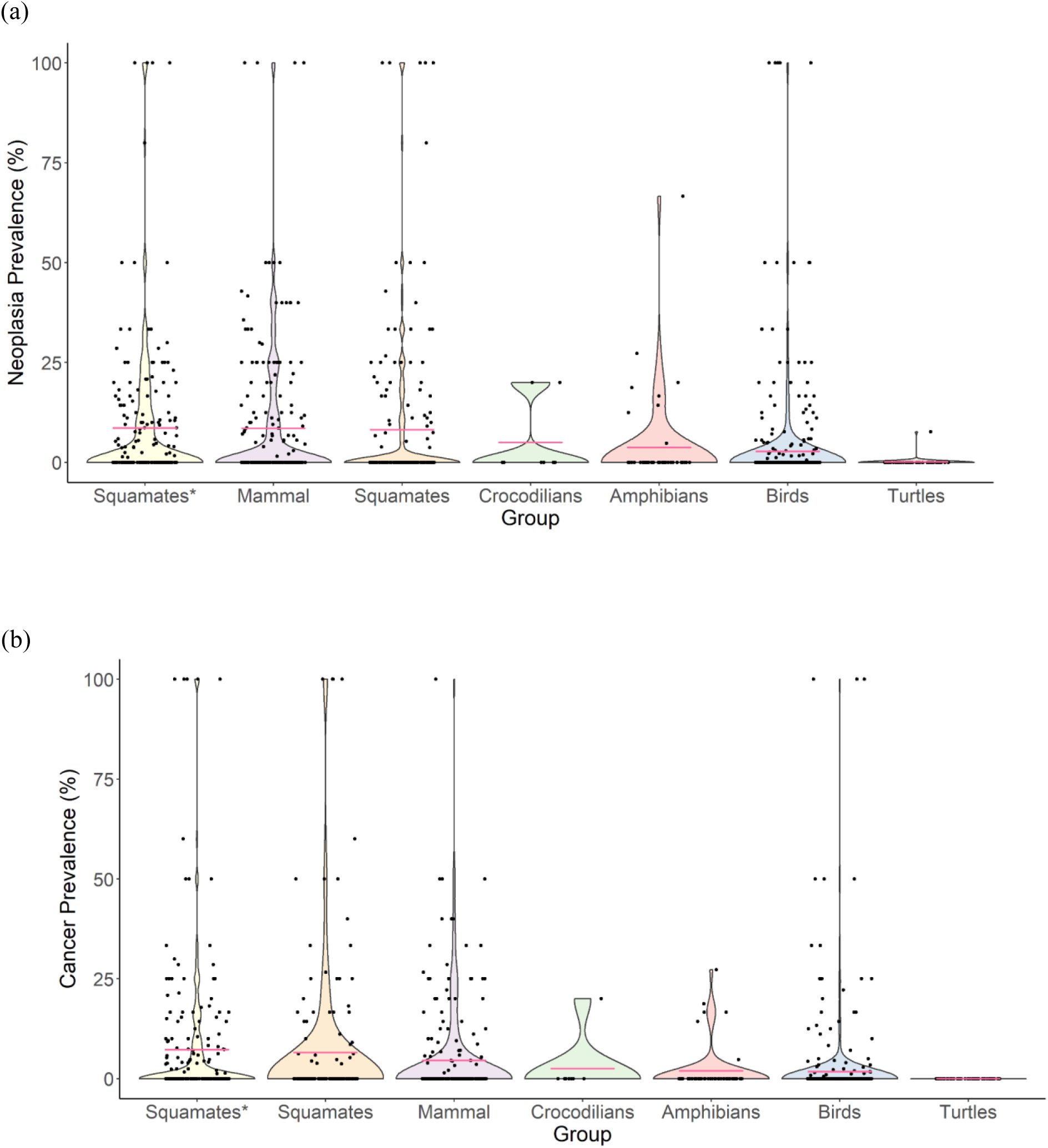
Summary of (a) neoplasia prevalence and (b) cancer prevalence across tetrapod groups. Each point represents the prevalence of a single species. Pink lines represent the mean prevalence of each group. Groups ordered left to right by descending mean prevalence. Prevalence is based on data as in Table 1. *Indicates squamate data from ^18^ included.

**Figure 2.**
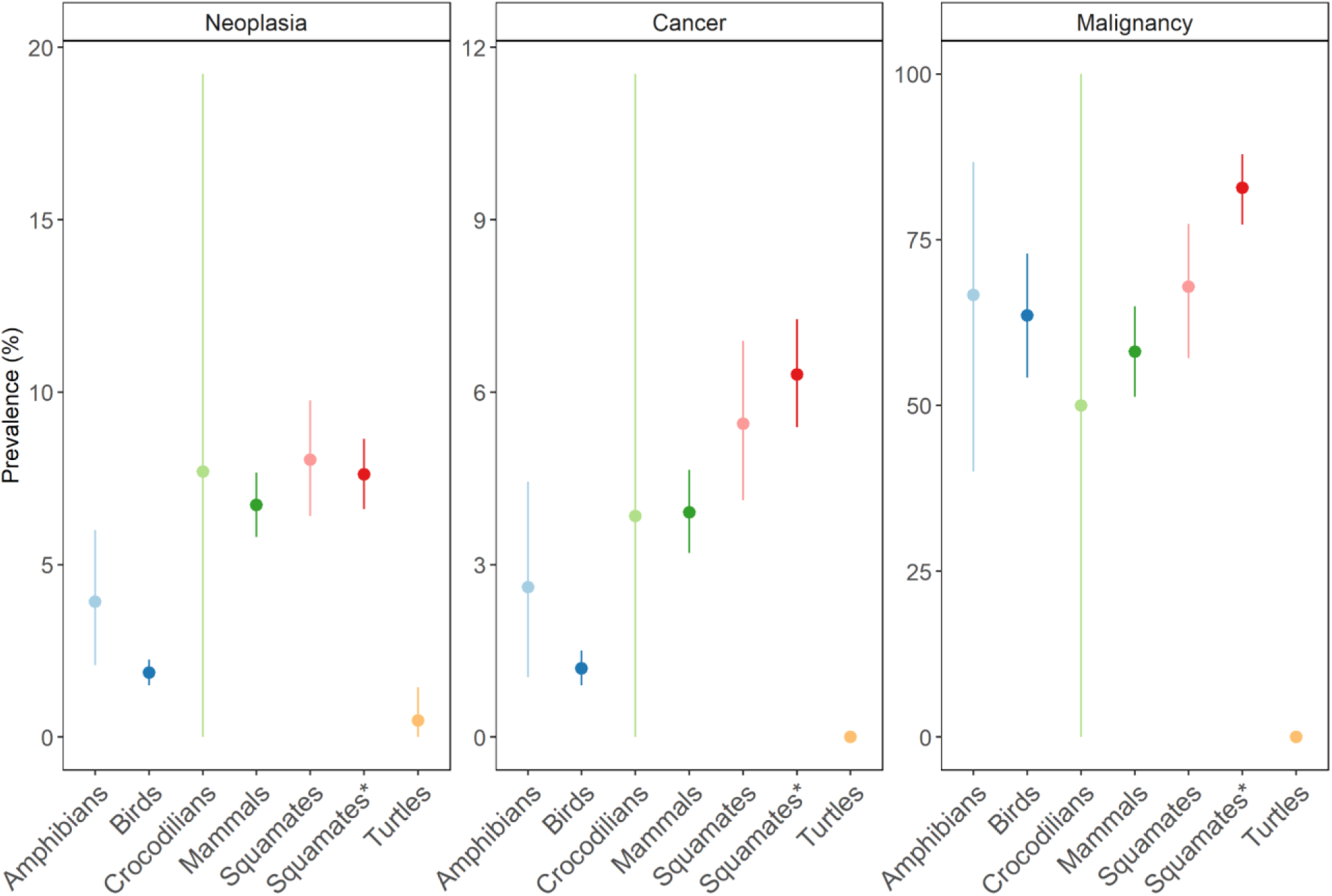
Prevalence of neoplasia, cancer, and malignancy transformation rate across tetrapod groups using the full dataset as in **Table 1**. Malignancy transformation rates is indicated as “malignancy prevalence” and was calculated as neoplasia count divided by cancer count. Points represent observed prevalence values and lines represent the 95% confidence intervals surrounding the observed prevalence value. *Indicates squamate dataset with data from ^18^ included. Please notice that due to the large variation in percentage among neoplasia, cancer, and malignancy transformation rate, the scale of the y-axis is different in the three graphs.

**Figure 3.**
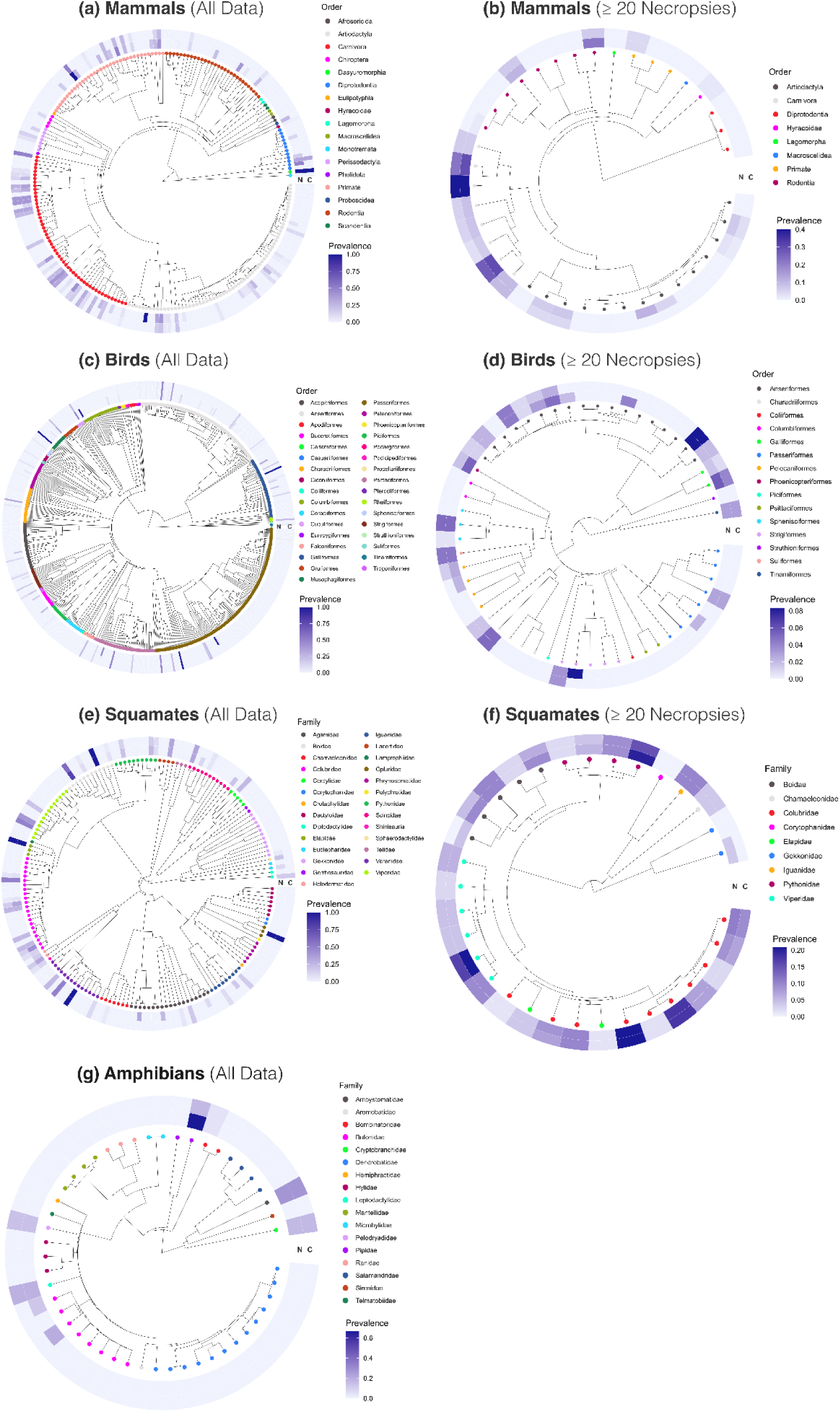
Interspecific variation in neoplasia (N) and cancer (C) prevalence within a) mammals including all species, b) mammals including only species with 20 or more necropsies, c) birds including all species, d) birds including only species with 20 or more necropsies, e) squamates, including all species, f) squamates including only species with 20 or more necropsies, and g) amphibians including all species. Taxa are color coded either by order (mammals and birds) or family (squamates and amphibians).

Amphibians, birds, mammals, and squamates had outlier species with prevalence values (neoplasia, cancer, or malignancy rate) significantly different (p<0.05) from the group mean (**Supplementary Materials Table 3**). Notably, among the species that were different from the group mean and a sample size of at least 20 necropsies per species, some outliers for neoplasia prevalence were the domestic chicken (7/157, 5.6%), the common hamster (24/114, 22%), and the diamondback rattlesnake (6/21, 29%). Additionally, three mammal outliers had 0% neoplasia prevalence despite a large number of necropsies per species: the artiodactyl collard peccary (0/89), domestic goat (0/91), and the rodent Patagonian mara (0/103).

Within our dataset, neoplasia prevalence ranked from highest to lowest as follows: 8% in both squamates and crocodilians, 7% in mammals, 4% in amphibians, 2% in birds, and 0.5% in turtles (**Table 1** and **Figure 2**). Cancer prevalence ranked from highest to lowest as follows: 5% in squamates, 4% in both mammals and crocodilians, 3% in amphibians, 1% in birds, and 0% in turtles. Malignancy transformation rate (i.e., cancer divided by neoplasia) was highest in squamates and amphibians with both groups having 70% malignancy transformation rate (80% in squamates when including data from ^18^), followed by mammals and birds with 60%, and crocodilians with 50% (**Table 1** and **Figure 2**). Squamates continue to have the highest neoplasia prevalence and malignancy transformation rate at ≥10 and ≥20 necropsies per species (S**upplementary Materials Table 4**). The observed prevalence values and their 95% confidence intervals among tetrapod groups indicated that birds and turtles had much lower neoplasia and cancer prevalence than other groups, but birds were similar to other groups for malignancy transformation rate (**Figure 2**). Except for crocodilian and amphibian cancer and malignancy transformation rates, 95% confidence intervals for prevalence values were narrow (**Figure 2**), indicating that estimates were precise.

Pairwise comparisons of neoplasia, cancer, and malignancy transformation rate calculated using *prop.test* and *p.adjust* with the BH method to account for multiple testing further supported differences among tetrapod groups (**Table 2**). Birds and turtles, with the lowest neoplasia and cancer prevalence, were not significantly different from each other but differed from all other groups (except crocodilians, **Table 2**). For neoplasia, amphibians differed from squamates, which had the highest neoplasia prevalence. Amphibians and squamates also differed from each other for cancer prevalence, but not for malignancy transformation rate (**Table 2**). Finally, mammals and squamates did not differ from each other in terms of neoplasia prevalence, but they were found to be different for cancer and malignancy transformation rate (**Table 2**). These trends were generally supported when the data was curated to only species with ≥10 necropsies (**Supplementary Materials Tables 5**). Birds differed from mammals and squamates for neoplasia and cancer for ≥20 necropsies, while only squamates differed from the other two groups for malignancy transformation rate (**Supplementary Materials Tables 6**).

**Table 2:** Pairwise comparison results for (a) neoplasia, (b) cancer, and (c) malignancy prevalence proportions run on full dataset. (as in **Table 1) using *prop.test*** a*nd **p.adjust*** with the Benjamini & Hochberg (“BH”) method (R package {stats}). Numbers in parentheses are proportion numbers per group calculated as # group neoplasia / # group total necropsy or # group cancer/ # group necropsy total, and # group cancer / # group total neoplasia for malignancy. Malignancy transformation rates is indicated as “malignancy prevalence” and was calculated as neoplasia count divided by cancer count. Significant values are in bold. Malignancy transformation rate could not be calculated for turtles since there were no instances of cancer in these groups. The asterisk denotes squamates data included from ^18^.

| GROUP NEOPLASIA PREVALENCE |  |  |  |  |  |  |  |
| --- | --- | --- | --- | --- | --- | --- | --- |
|  | Amphibians<br>(15/383) | Birds<br>(96/5124) | Crocodilian<br>s (2/25) | Mammals<br>(191/2841) | Squamates<br>(84/1053) | Squamates*<br>(198/2608) | Turtles<br>(1/207) |
| Turtles<br>(1/207) | 0.07 | 0.3 | 0.07 | **2.70E-03** | **1.50E-15** | **1.10E-03** | 1 |
| Squamates*<br>(198/2608) | **0.03** | **1.50E-15** | 1 | 0.3 | 1 | 1 |  |
| Squamates<br>(84/1053) | **0.03** | **1.50E-15** | 1 | 0.3 | 1 |  |  |
| Mammals<br>(191/2841) | 0.1 | **1.50E-15** | 1 | 1 |  |  |  |
| Crocodilian<br>s (2/25) | 1 | 0.2 | 1 |  |  |  |  |
| Birds<br>(96/5124) | **0.03** | 1 |  |  |  |  |  |

(b)
| GROUP CANCER PREVALENCE |  |  |  |  |  |  |  |
| --- | --- | --- | --- | --- | --- | --- | --- |
|  | Amphibians<br>(10/383) | Birds<br>(61/5124) | Crocodilian<br>s (1/25) | Mammals<br>(111/2841) | Squamates<br>(57/1053) | Squamates*<br>(164/2608) | Turtles<br>(0/207) |
| Turtles<br>(0/207) | **1.50E-15** | 0.4 | 0.4 | **0.02** | **0.004** | **0.002** | 1 |
| Squamates*<br>(164/2608) | **0.02** | **1.50E-15** | 1 | **4.0E-4** | 0.7 | 1 |  |
| Squamates<br>(57/1053) | 0.1 | **1.50E-15** | 1 | 0.1 | 1 |  |  |
| Mammals<br>(111/2841) | 0.5 | **1.50E-15** | 1 | 1 |  |  |  |
| Crocodilians<br>(1/25) | 1 | 1 | 1 |  |  |  |  |
| Birds<br>(61/5124) | 0.08 | 1 |  |  |  |  |  |

(c)
| **GROUP MALIGNANCY PREVALENCE** |  |  |  |  |  |  |
| --- | --- | --- | --- | --- | --- | --- |
|  | Amphibians<br>(10/15) | Birds<br>(55/82) | Crocodilians<br>(1/2) | Mammals<br>(106/185) | Squamates<br>(50/75) | Squamates*<br>(140/173) |
| Squamates*<br>(140/173) | 0.8 | **0.004** | 1 | **3.20E-06** | **0.05** | 1 |
| Squamates<br>(50/75) | 1 | 1 | 1 | 0.8 | 1 |  |
| Mammals<br>(106/185) | 1 | 1 | 1 | 1 |  |  |
| Crocodilians<br>(1/2) | 1 | 1 | 1 |  |  |  |
| Birds<br>(55/82) | 1 | 1 |  |  |  |  |

Within each studied vertebrate group, we found wide variation in neoplasia and cancer prevalence among families and orders (**Figure 3, Supplementary Materials Table 7,** full comparisons on Dryad *after manuscript acceptance*).

### Intrinsic Cancer Risk Versus Neoplasia, Cancer, and Malignancy Transformation Rate

Intrinsic cancer risk corresponds to the predicted risk of developing cancer based on a combination of species lifespan and body mass ^20,21^. Across tetrapods, we found ICR to be highly variable (**Figure 4**). Certain species showed extraordinarily high ICR values compared to other species in their groups, including Chinese giant salamanders (*Andrias davidianus*), common ostriches (*Struthio camelus*), Asian elephants (*Elaphas maximus*), Komodo dragons (*Varanus komodoensis*), and Galapagos tortoises (*Geochelone nigra* complex). Overall, based on the species in our dataset, amphibians and turtles had lower average ICR values than other tetrapod groups (**Figure 4**). Note that the ICR calculation (ICR = lifespan^6^ x body mass) is more strongly affected by variation in lifespan than body mass.

**Figure 4.**
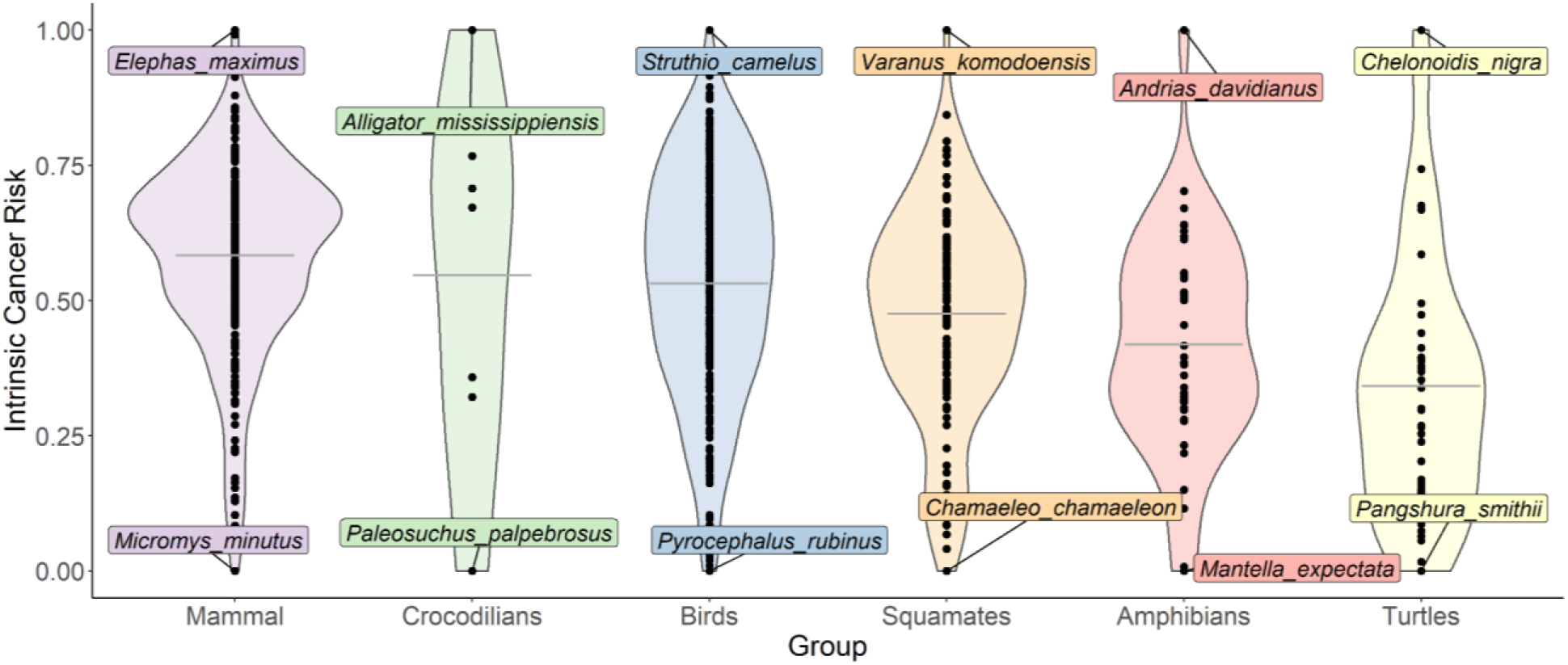
Relative estimated intrinsic cancer rates (ICR) based on body mass and lifespan data for all tetrapod species examined in this study (see Materials and Methods for ICR calculation). Gray lines in each violin represent the mean ICR for the tetrapod group. For visual representation, original ICR values were log scaled and transformed using a min-max normalization (ICR – min(ICR))/(max(ICR) – min(ICR)) to bring them to a 0-1 range within each group, where 0 and 1 represent the species with the lowest and highest intrinsic cancer risk, respectively, within each lineage. Please note that species with 1 in the graph do not necessarily have a 100% intrinsic cancer rate, but they just correspond to the higher intrinsic rate in the group.

Using phylogenetic comparative methods (*gee*), we found significant relationships between ICR and neoplasia prevalence in squamates (positive, p=0.007) and mammals (negative, p=0.0002), and between ICR and cancer prevalence in amphibians (positive, p=0.002; **Table 3**). ICR was a highly correlated to neoplasia or cancer prevalence for squamates, mammals, and amphibians, but not for other groups, and was not related to malignancy transformation rates in any group.

**Table 3.**
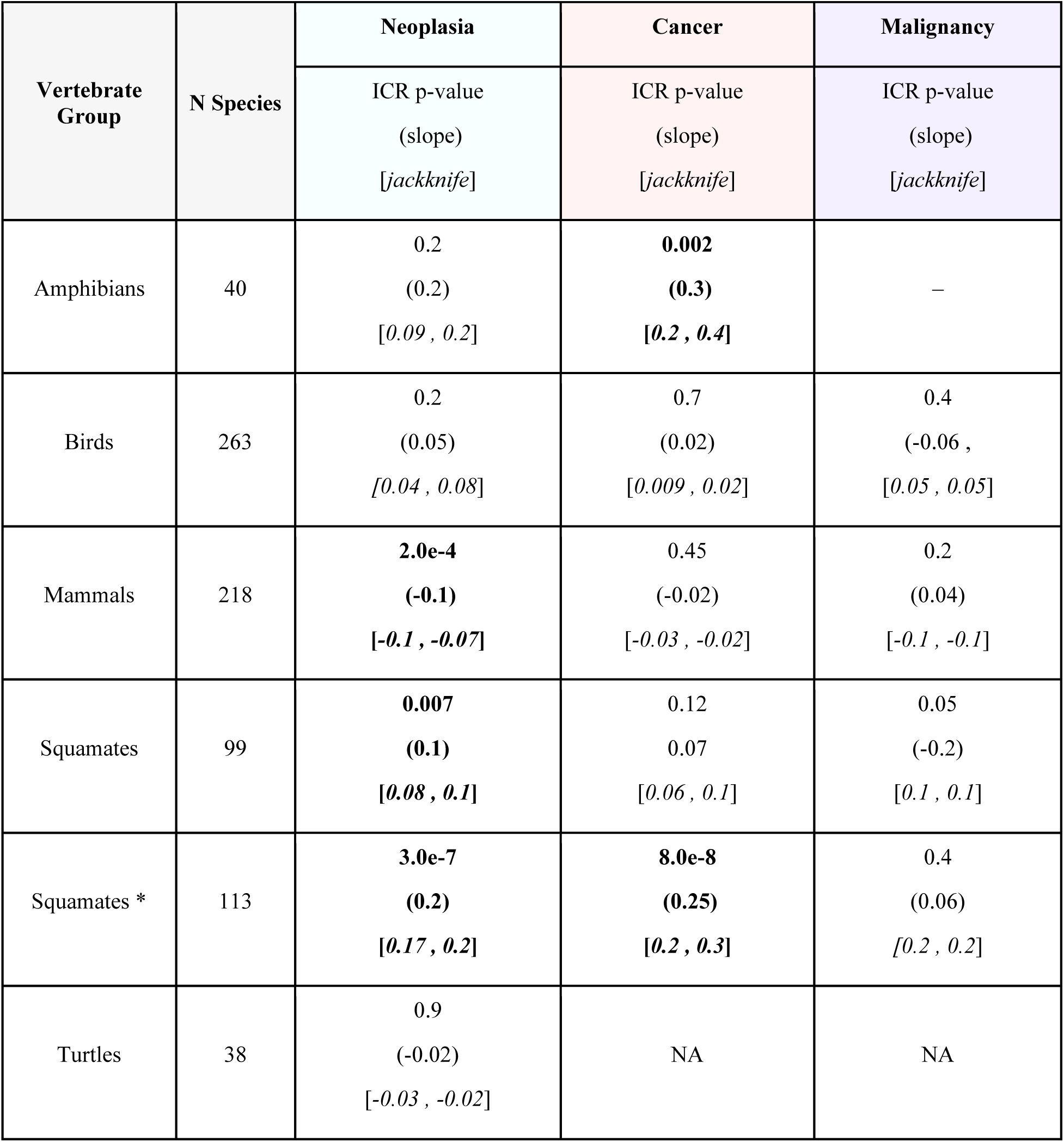
Influence of intrinsic cancer risk (lifespan^6^ x body mass) on neoplasia, cancer or malignancy transformation rate. Malignancy transformation rate is indicated as “Malignancy” in the table. Analyses were run using the generalized estimating equations (*gee*; see Materials and Methods for additional information). Species included in analysis were curated to those for which both lifespan and body mass life history traits were available. P-values <0.05 are in bold with slope coefficients in parentheses to show direction effect and jackknife 95% confidence intervals italicized in brackets. Dashed lines represent models that did not run due to non-converging. NA in turtles indicates that analyses could not be run because cancer was not observed in these groups.

### Neoplasia, Cancer, and Malignancy Transformation Rate as a Function of Body Mass and Lifespan

We also tested the influence of body mass or lifespan separately on the prevalence of neoplasia, cancer, and malignancy transformation rates in each tetrapod group, while accounting for phylogenetic relationships among species within each group. We found that body mass was positively correlated with neoplasia and cancer in amphibians, neoplasia in squamates, and negatively correlated to neoplasia and malignancy transformation rate in mammals (**Table 4**). Lifespan was positively correlated with cancer in amphibians, neoplasia and cancer in squamates, and malignancy transformation rates in mammals, and negatively correlated with neoplasia in mammals (**Table 4**). Thus, in amphibians and no other studied group, both body mass and lifespan influence cancer, while in mammals and no other group, both body mass and lifespan influence neoplasia and malignancy transformation rate (**Table 4**). A positive correlation means that when taking into account phylogenetic relationships among the studied species, an increase in that life history trait correspond to higher prevalence, which supports our predictions using ICR (**Table 3**). On the other hand, in mammals, we found a negative relationship between body mass and lifespan and neoplasia, as well as body mass and malignancy transformation rates, meaning that increases in that life history trait correspond to a decrease in prevalence (**Table 4**). For groups with ≥20 species and ≥10 or 20 necropsies per species (birds, mammals, and squamates), lifespan influenced neoplasia and malignancy transformation rate in mammals, and neoplasia and cancer in squamates (**Supplementary Materials Table 8**). To note that while a threshold of 10 or 20 or more necropsies per species has been used in previous studies, due to the reduced species representation and prevalence distribution on each of the group phylogenies (**Figure 3**), phylogenetic heterogeneity in the data is lost and analyses fail to converge due to lack of variation in the data across the reduced phylogenetic representation. The fact that less tests reach convergence or are significant when species with few individuals are dropped from the analyses it is further illustrated by the jackknife confidence intervals. 95% confidence intervals in fact increase when the number of species included in the analysis is reduced (see confidence intervals of **Table 4 and Supplementary Materials Table 8**) The meaning of this is a reduced confidence in the results obtained using less species with more necropsies than the entire dataset.

**Table 4.** Influence of body mass and lifespan on cancer, neoplasia, and malignancy prevalence. Analyses run using the full dataset on cancer, neoplasia, and malignancy transformation rates using either lifespan or body mass as the predicting variable and taking into account group’s phylogenetic relationships. Analyses were run using the generalized estimating equations (*gee*; see Materials and Methods for additional information). Malignancy transformation rates is indicated as “malignancy prevalence” and was calculated as neoplasia count divided by cancer count. P-values <0.05 are in bold with slope parameters in parentheses to show direction effect and jackknife 95% confidence intervals italicized in brackets. Lines represent models that did not run due to non-converging (see Materials and Methods for additional information). Numbers in the column “N Species” refers to the number of species per trait being analyzed. If a species mass or lifespan history trait was unknown it was removed from that analysis. Crocodilians could not be analyzed due to an insufficient number of species. Turtles could not be analyzed for cancer and malignancy transformation rate as no instances of cancer were found.

| Vertebrate Group | Total Neoplasia |  |  |  | Cancer |  |  |  | Malignancy |  |  |  |
| --- | --- | --- | --- | --- | --- | --- | --- | --- | --- | --- | --- | --- |
|  | Mass |  | Lifespan |  | Mass |  | Lifespan |  | Mass |  | Lifespan |  |
|  | N Species | p-value (slope) [jackknife] | N Species | p-value (slope) [jackknife] | N Species | p-value (slope) [jackknife] | N Species | p-value (slope) [jackknife] | N Species | p-value (slope) [jackknife] | N Species | p-value (slope) [jackknife] |
| Amphibians | 47 | **0.003**<br>(1.6)<br>[1.3, 2.1] | 42 | 0.31<br>(0.7)<br>[0.2, 1.5] | 47 | **0.0014**<br>(1.5)<br>[1.3, 2.3] | 42 | **0.009**<br>(1.8)<br>[1.1, 2.2] | 8 | 0.85<br>(0.3)<br>[0.12, 1.4] | 8 | — |
| Birds | 299 | 0.4<br>(0.07)<br>[0.07, 0.1] | 267 | 0.3<br>(0.3)<br>[0.3, 0.4] | 299 | 0.6<br>(0.07)<br>[0.06, 0.08] | 267 | 0.96<br>(-0.02)<br>[0.03, 0.08] | 57 | 0.12<br>(0.3)<br>[0.2, 0.3] | 48 | 0.3<br>(-0.5)<br>[-0.7, -0.4] |
| Mammals | 220 | **0.002**<br>(-0.2)<br>[-0.2, -0.15] | 218 | **0.001**<br>(-0.8)<br>[-0.8, -0.4] | 220 | 0.6<br>(0.03)<br>[-9e-4, 0.05] | 218 | 0.3<br>(-0.3)<br>[-0.4, -0.3] | 77 | **0.005**<br>(-0.2)<br>[-0.3, -0.2] | 77 | **0.001**<br>(1.1)<br>[1.0, 1.1] |
| Squamates | 102 | **0.02**<br>(0.2)<br>[0.16, 0.2] | 122 | 0.2<br>(0.4)<br>[0.3, 0.6] | 102 | 0.14<br>(0.1)<br>[0.09, 0.2] | 122 | 0.8<br>(0.08)<br>[-0.02, 0.16] | 32 | 0.07<br>(-0.4)<br>[0.4, -0.2] | 32 | 0.2<br>(-0.9)<br>[-1.1, -0.7] |
| Squamates * | 116 | **0.0003**<br>(0.24)<br>[0.2, 0.3] | 142 | **0.0003**<br>(0.9)<br>[0.8, 1.1] | 116 | **0.0004**<br>(0.3)<br>[0.2, 0.3] | 142 | **0.0004**<br>(1.07)<br>[1.0, 1.2] | 56 | 0.6<br>(0.1)<br>[0.05, 0.2] | 56 | 0.5<br>(0.4)<br>[0.07, 0.6] |
|  | Mass |  | Lifespan |  | Mass |  | Lifespan |  | Mass |  | Lifespan |  |
|  | N Species | p-value (slope) [jack knife] | N Species | p-value (slope) [jack knife] | N Species | p-value (slope) [jack knife] | N Species | p-value (slope) [jack knife] | N Species | p-value (slope) [jack knife] | N Species | p-value (slope) [jack knife] |
| Turtles | 46 | 0.8 (0.07) [0.05, 0.07] | 42 | 0.8 (-0.3) [-0.6, 0.13] | NA | NA | NA | NA | NA | NA | NA | NA |

\*Includes data from <sup>18</sup>

### Re-analysis of Publicly Available Data from Previous Studies on Mammals

Previous studies in mammals ^3,4^ have found that change of body mass and lifespan is not associated with a change of neoplasia and cancer prevalence (but see ^10^ on the influence of body weight after correcting for gestational period). To assess the influence of our statistical approach on the results obtained from prior studies, we re-analyzed their data using *gee*. Based on the data from Boddy et al. ^3^, which included 36 species (28 of them with ≥10 necropsies per species), we found no influence of body mass or lifespan change on neoplasia or cancer prevalence evolution, supporting their results (**Supplementary Materials Table 9**). However, as their interspecific sampling is lower than ours and as parameter estimates were close to 0 (**Supplementary Materials Table 9**, see jackknife values), it could also mean that their sample size is too low to detect any relationships. However, when we used the data from Vincze et al. ^4^, including 189 species with 20 or more necropsies per species, our statistical approach indicated that both changes in body mass and life expectancy were significantly (p<0.001) related to changes in neoplasia prevalence, in contrast to what was found by the authors. Note that in Vincze et al. ^4^, the neoplasm was defined as the cause of death, so in this sense it is cancer mortality data and does not exactly correspond to any of the categories – neoplasia, cancer, or malignancy transformation rate – used in our study. The fact that we found a significant relationship between life history traits and cancer mortality data when using their dataset supports the relationship in mammals between cancer mortality and life history traits (**Supplementary Materials Table 9**).

## DISCUSSION

We estimated neoplasia and cancer prevalence and malignancy transformation rates across vertebrates, including amphibians, birds, crocodilians, mammals, squamates, and turtles, focusing on variation both within and among groups. Previous studies have explored cancer prevalence in birds and reptiles as a single taxon (sauropsids) ^5,10^ or focusing on birds alone ^6,13^, which limits understanding of cancer diversity among these groups. Our study addresses this gap by examining each vertebrate group individually. We found that neoplasia and cancer prevalence, as well as malignancy transformation rates, were higher in squamates, especially in large-bodied lizards like Helodermatidae (e.g., gila monster). This could be due to the greater number of cells associated with larger body sizes, increasing cancer risk. We also found high malignancy transformation rates almost uniformly across snake families (67-100%). In contrast, we found no cancer and only one case of neoplasia (*Chelus fimbriata*) in turtles. In crocodilians, only two out of eight species had neoplasia, one of which was malignant (*Osteolaemus tetraspis*).

Despite the remarkably low prevalence values, neoplasia and cancer do occur in turtles and crocodilians ^12,15,16,22,23^. These low rates may be due to slower metabolic rates, reduced oxidative stress, and other unique physiological traits. Previous work, which did not distinguish between benign and malignant tumors, found 1.2-2.7% neoplasia prevalence in turtles ^15,16^ and 2.2% in crocodilians ^15^. Turtles have lower germline mutation rates in mitochondrial and nuclear DNA than mammals and birds ^24^, possibly related to lower somatic mutation rates and lower cancer incidence. In addition, turtles were found to have slower aging rates and derived cellular phenotypes associated with delayed aging and cancer resistance ^25–27^.

We identified low rates of neoplasia and cancer in birds compared to mammals and reptiles, consistent with previous studies ^11,12^. This low prevalence may be related to the bursa of Fabricius, a specialized organ producing B-cells in juvenile birds, and enhanced immune system development^13^. Higher prevalence of neoplasia and malignancy transformation rate in some avian species has been correlated with larger clutch sizes ^6^. Our data confirmed that birds have very low neoplasia rates, but malignancy transformation rates are comparable to mammals and squamates. This might be due to the efficiency of their immune system in preventing neoplasia, but once tumors develop, they rapidly progress to malignancy. Although this effect could be due to bias in the types of tissues sent for pathology analysis, if confirmed, these results indicate that when neoplasia occurs in birds, it is usually malignant (>50% of the time).

Previous studies have found conflicting results regarding the influence of life history traits, such as body mass and lifespan, on neoplasia and cancer prevalence across tetrapod groups. Our work is the first to compare expected rates of intrinsic cancer risk (ICR) based on body mass and lifespan with observed rates across each studied vertebrate group. We found higher ICR to be correlated with higher neoplasia prevalence in squamates. Conversely, in mammals, higher ICR values correlate with lower neoplasia prevalence, possibly due to evolved tumor suppression mechanisms in larger, longer-lived species. ICR is also a good positive predictor of cancer in amphibians. However, ICR does not correlates to neoplasia or cancer in birds, turtles, or crocodilians, nor malignancy transformation rates across any studied vertebrate groups. These results highlight the complexity of cancer evolution across different vertebrate groups and underscores the need for taxon-specific approaches in future comparative oncology research.

When analyzing the influence of body mass and lifespan separately, at least one of these two life history traits has an influence on neoplasia or cancer in several vertebrate groups including amphibians, squamates, and mammals. Our results contradict a previous study that observed a positive association between body mass and neoplasia prevalence in birds ^13^. In mammals, change in cancer prevalence was previously found to be unrelated to body mass or lifespan evolution (but see ^10^), but instead potentially influenced by diet, with higher prevalence values for cancer observed in carnivores ^4,12^ or by gestation period and litter size ^3,10^. In contrast, we found a positive relationship between lifespan and malignancy transformation rate in mammals, but inverse relationships between body mass and both neoplasia and malignancy transformation rates as well as an inverse relationship between lifespan and neoplasia. This suggests that animals with larger body mass have lower neoplasia and malignancy transformation rates, and as such, larger animals have effectively evolved mechanisms to reduce mortality due to cancer by experiencing a decreased occurrence of neoplasia than except for their body size. At the same time, our results indicate a more complex relationship existing between lifespan, neoplasia, and malignancy transformation. Longer-lived species may have evolved mechanisms protecting them from developing neoplasia, but once it occurs, it may often become malignant, hence the positive relationship with malignancy transformation.

In this work, we have established the novel findings that squamates and amphibians both have a positive relationship between body mass and neoplasia prevalence, and amphibians also have a positive relationship between body mass, lifespan, and cancer. It has been proposed that regenerative capacities, as observed in some amphibian species, may be associated with tumor suppression mechanisms ^28,29^, and that regeneration in amphibians, unlike mammals, does not necessarily result in increased cancer prevalence ^30,31^. This hypothesis needs further exploration since we found that amphibians had lower neoplasia and cancer prevalence than mammals and squamates, but higher than birds. Regenerative capacities are not uniform across species, orders, and life-stages of amphibians ^32^, possibly explaining why neoplasia and cancer prevalence is relatively high in this group. Finally, we found that neoplasia, cancer, and malignancy transformation rates are remarkably frequent in snakes and occurred in almost all snake families included in this study, which is similar to ^18^.

Our study pioneers an approach for analyzing the relationship between life history traits and cancer prevalence data, marking a significant advancement in the field. Unlike previous studies that treated cancer prevalence as a continuous variable using Phylogenetic Generalized Least Squares (PGLS) for correlation analysis ^3,4^, we recognize that prevalence data are proportional, adhering to a binomial rather than a normal distribution ^17^. The skewness towards zero in cancer occurrence across species challenges the PGLS method’s assumption of a normal distribution. We employ Generalized Estimating Equations (*gee*) to encompass the binomial distribution and account for the skewed nature of cancer prevalence data. This method allows for the inclusion of all collected data, improving representation even for species with sparse samples. *Gee* utilizes individual neoplasia and cancer counts instead of averaging prevalence per species, enabling a more detailed analysis that considers variations in sample size and distribution across the phylogeny. Our analysis, conducted without imposing a minimum threshold for necropsies per species, and then with subsets of species having at least 10 or 20 necropsies, underscores the limitations of data censoring. Excluding species based on arbitrary necropsy thresholds reduces phylogenetic representation and variation in prevalence across clades, potentially leading to non-converging analyses or insignificantly large confidence errors. Thus, we advocate for using statistical methods suited for proportional data and individual rather than mean species data.

Finally, to assess whether our analytical approach may explain why our findings differ from previously published work, we reanalyzed the mammalian data from both Boddy et al. ^3^ and Vincze et al. ^4^ using our statistical methodology. We found that when the Vincze et al.’s ^4^ data were reanalyzed with our approach, their data did support a relationship between cancer mortality and life history traits. This suggests that a relationship between life history traits and neoplasia or cancer mortality may previously not have been recovered due to the use of less suitable statistical approaches.

### Caveats And Future Directions

Our study has identified a significant influence of body mass and lifespan on neoplasia and cancer prevalence across various tetrapod groups. The additional jackknife analysis we conducted suggests that limitations such as sparse data, long evolutionary branches within the phylogeny of focal species, and the uneven distribution of neoplasia, cancer, or rates of malignancy transformation across the phylogeny can affect the outcomes of comparative phylogenetic methods. Addressing these limitations by expanding the dataset both in terms of species diversity and the number of necropsies per species could mitigate some of these issues. However, the acquisition of more data is challenged by the fact that certain species, especially those in zoos like turtles, are endangered or critically endangered ^33^, limiting the availability of individuals for study. Analyzing data from all available individuals ensures a more comprehensive phylogenetic representation and strengthens the robustness of the results.

Acknowledging potential biases is crucial in studies of this nature. The frequency and type of tissue samples submitted for pathological analysis can vary significantly across vertebrate groups, zoos, and over time, influenced by evolving necropsy practices. A strategy to minimize such biases involves comprehensive sampling across a wide range of groups, zoological institutions, and geographical areas. Furthermore, the zoo environment, predominantly housing captive-bred species often at risk of inbreeding, may influence cancer patterns differently than in wild populations. Although zoo data provide invaluable insights for answering broad comparative questions, as demonstrated in our study, future research should aim to address these considerations to enhance the reliability and applicability of findings. Finally, since animals housed in zoos may have different lifespans and aging processes compared to their wild counterparts ^34^, obtaining data on cancer prevalence in wild animals is essential for a complete understanding of cancer occurrence in these organisms. However, accurately measuring cancer prevalence in wild animals is challenging, especially for elusive species. Sick or dead animals are likely to be eaten or difficult to find, leading to significant underestimates of cancer prevalence.

### Conclusions

Our findings reveal that vertebrate groups offer distinct insights into the evolution of cancer suppression mechanisms. In mammals, our data support Peto’s Paradox, showing that evolutionary pressures have favored the suppression of neoplasia in larger species. We also observed a positive correlation between malignancy transformation rates and lifespan in mammals, suggesting that neoplasia, once formed, are prone to becoming malignant. This pattern implies that evolutionary strategies may have prioritized preventing neoplasia over cancer, guiding future research toward uncovering specific cancer suppression mechanisms in mammals, including humans.

In amphibians and squamates, body mass and lifespan emerged as reliable predictors of neoplasia prevalence, although this did not extend to malignancy transformation rates. In contrast, birds exhibited relatively low rates of both neoplasia and cancer, hinting at evolved resistance mechanisms to neoplasia irrespective of their life history traits. However, the high rate of malignancy transformation in birds suggests that neoplasia that do develop are likely to become malignant.

Our study also highlights the urgent need for more comprehensive data on amphibians, a group where the dual influence of lifespan and body mass on cancer prevalence was noted, yet remains underrepresented in comparative oncology research. The limited species representation in our dataset indicates a gap in understanding cancer across vertebrates.

Turtles, and possibly crocodilians, stand out as valuable models for cancer resistance research due to their exceptionally low neoplasia rates and even rarer malignancy transformation. These findings underscore the diversity of cancer prevalence and resistance mechanisms across vertebrate groups and emphasize the importance of broadening the scope of comparative oncology to include a wider array of species.

## MATERIALS AND METHODS

### Cancer Data Collection and Curation

We searched paper reports or digital databases of six European zoos (Allwetterzoo, Münster, Germany; ZOOM Erlebniswelt Gelsenkirchen, Gelsenkirchen, Germany; Der Grüner Zoo Wuppertal, Wuppertal, Germany; Wildlife Zoo Hellenthal, Hellenthal, Germany; Rotterdam Zoo, Rotterdam, The Netherlands; Zoo Zürich, Zürich, Switzerland) and one in the United States (Birmingham Zoo, Alabama) for necropsies recorded between 1998 and 2019 (**Supplementary Material Table 1**). The databases for Allwetterzoo, ZOOM Erlebniswelt, Zoo Wuppertal, Zoo Hellenthal, Zürich Zoo, and Birmingham Zoo were all local databases, while Rotterdam Zoo data was obtained through local paper reports and the Species360 Zoological Information Management System (ZIMS ^35^) with Rotterdam Zoo’s authorization. We also used a dataset of neoplasia (benign or malignant) in snakes from Duke et al. ^18^, which is based on cancer prevalence calculated from the number of biopsies and necropsies of the total number of individuals (live and dead) of each species housed at a given time at participating institutions in the study. Since this prevalence was partly based on live individuals instead of strictly necropsy reports, like the rest of our data, we performed subsequent analyses both with and without the data from Duke et al. ^18^.

The list of keywords used to search for neoplasia within the necropsy reports can be found in the file “Neoplasia_Terms” (**Supplementary Materials**). To build the dataset for this study, we only included tumor data confirmed by histological reports from veterinary pathologists. Any neoplastic individual that was too autolytic to diagnose potential cancer incidence was removed from the study altogether. To be conservative in our dataset, any individual that died during the first month of life was removed from the dataset since they may have a low risk of developing cancer, which would bias the data toward lower cancer prevalence. To note that although the data in this study all come from necropsy reports (except from ^18^), the cause of death in these reports was not necessarily recognized to be the tumor(s).

We tallied the total number of necropsies, independently of whether they had a tumor or not, for each species represented by a necropsy report during the time frame considered in this study (1998-2019). The data were organized into cases of ‘neoplasia’ (any benign or malignant neoplasia confirmed by histology) and ‘cancer’ (malignant neoplasia diagnosed by abnormal cellular nuclei, tissue invasion, and/or metastasis). Finally, two species, *Rousettus aegyptiacus* (mammal) and *Thamnophis radix* (squamate from ^18^) had more than 500 necropsies and were removed from the dataset as outliers, defined as having necropsies more than three standard deviations above the group mean.

“Neoplasia prevalence” per group was calculated as the total number of neoplasias divided by the total number of necropsies for each group following the rate of tumorigenesis calculation in ^36^. “Cancer prevalence” per group was calculated as the total number of cancers divided by the total number of necropsies for each group. “Malignancy transformation rate” per group was calculated as the total number of cancers divided by the total number of neoplasias for each group also following ^36^. Finally, although malignancy requires neoplasia to occur first ^36^, our data are based only on deceased individuals (i.e., necropsies). Thus, the number of neoplasias could be skewed if the individual did not die from it or other causes. For that reason, we report neoplasia, cancer, and malignancy transformation rate values throughout our results. The full dataset can be found on Dryad (*after manuscript acceptance*).

### Life History Data Collection

Maximum body mass and lifespan were collected for each species when available using the Animal Aging and Longevity Database (AnAge) ^37^. If no species data was found in AnAge, primary sources were used when available. Data origin for life history trait information is listed in the full dataset file under “Sources LHT” for each group (*Dryad after manuscript acceptance*). If no source for a verified maximum lifespan or body mass could be found, then that species was removed from life history trait analyses. Most amphibian life history trait data were collected from the AmphiBIO database ^38^. As body mass information was lacking for most amphibians, snout-vent length was used as a proxy for body mass. This is the most common measurement for amphibian body size since mass can be highly variable within the same species and within individuals, even over a short time period, due to factors such as hydration, bladder size, physiological state, and reproductive state ^39,40^.

### Intrinsic Cancer Risk Analysis

Intrinsic cancer risk (ICR) due to body size and lifespan was estimated for each species in the dataset following Peto’s model ^20^, where ICR = lifespan^6^×body mass. Only species for which both body mass and lifespan data are both available could be included in this analysis. To test the relationship between expected cancer risk and observed neoplasia, cancer, and malignancy transformation rate for each group, while taking phylogenetic relationships into account, we used generalized estimating equations (*compar.gee*, R package {ape}) ^17,41^ between the log scaled intrinsic cancer risk and neoplasia, cancer, or malignancy transformation rate for each group using R v4.1.1 ^42^.

### Statistical Analyses

The Shannon-Wiener Diversity Index and the Shannon Equitability Index were used to assess species diversity and the evenness of species distribution in our dataset ^43,44^. Additional information on Methods and Results for this can be found in the Supplementary Materials (**Supplementary Materials Table 2**).

To determine if the prevalence of neoplasia, cancer, and malignancy transformation rates differ significantly among groups, we calculated 95% confidence intervals for each group’s prevalence values separately. Prevalence of neoplasia, cancer and malignancy transformation rate among tetrapod groups were compared in pairwise fashion using the test of equal proportion using the R function *prop.test* and ***p.adjust*** with the Benjamini & Hochberg (“BH”) method to account for multiple testing (R package {stats}). To our knowledge, this is the first work that considers pairwise comparisons of cancer prevalence within and between each group, thus assessing which prevalence values are significantly different from one another. Between species, prevalence, using neoplasia/cancer incidence and sample size, were tested against the mean of their respective group using *prop.test* (R package {stats}) to identify outliers with significantly (p = <0.05) different prevalence from the group mean (**Supplementary Materials Table 3**). To assess the influence of lifespan or body mass on neoplasia, cancer, and malignancy transformation rate, statistical analyses were run for neoplasia prevalence, cancer prevalence, and malignancy transformation rate. Analyses were run for the entire dataset and also considering species with ≥10 and 20 necropsies per species – both thresholds used in previously published papers on this topic ^2–4,6,10^ (**Supplementary Materials Table 4** contains necropsy sample sizes for each group for which 20 or more species were available with 10 and 20 or more necropsies). We repeated analyses with the full dataset and with species having more necropsies to ensure higher phylogenetic resolution and avoid bias from overrepresented clades (**Figure 3**). Furthermore, as in the statistical approach used in this paper, individual data and not species prevalence are used for the analyses and the relative weight of the number of individuals per species is taken into account by this approach, we want to assess the influence of using a different number of necropsies per species on the results. Finally, in order to obtain an idea of the influence of interspecific sampling within groups, analyses were run by jackknifing species of the dataset and a 95% confidence interval was obtained for the estimated parameters.

For the statistical analyses, mass and lifespan data were log-transformed to linearize potential allometric effects. We used generalized estimating equations (*compar.gee*) to estimate the influence of body mass or lifespan on neoplasia, cancer, and malignancy transformation rate while considering the phylogenetic relationships within each group ^17,41^. As cancer prevalence is the response variable and is a proportion, the binomial family was used for the model and the number of positive (occurrence of neoplasia, cancer) and negative (non-occurrence) individuals was entered as raw data for each species. For these comparative analyses in which phylogenetic relationships and each life history trait was analyzed, we only used species that could be matched with a published phylogeny for their group and for which at least one life history trait (lifespan and/or body mass) could be found online or in literature (see Life History Data Collection). The novelty of this approach is that instead of using proportions for neoplasia, cancer, or malignancy transformation rate per species, as commonly done ^3,4,6,10^, we directly used individual count for each species. This ensures that per species sample size is considered directly in the model, without the need of giving different weights according to sampling; this also helps to directly model the prevalence as a logistic response. Numbers of species per group for all analyses can be found in **Table 1** and in the Supplementary Materials (**Table 4**). These analyses were not run for the crocodilians as there were too few species to allow convergence of the model.

Phylogenetic trees for each studied group were obtained from the following sources: amphibians ^45^, birds ^46^, crocodilians ^47^, mammals ^48^, squamates ^49^, and turtles ^50^. All phylogenetic trees were transformed to ultrametric trees using the *chronos* function in R package [ape], using default parameters (no fossil calibration). All the statistical analyses were run in R v4.1.1.

### Re-analysis of Previously Published Results on Mammals

To assess the influence of our statistical approach on the results obtained from previous studied, we reanalyzed data from Boddy et al. ^3^ and Vincze et al. ^4^ separately using *gee*. Body mass and lifespan data were log transformed as the authors did in their papers. For the Vincze et al. ^4^ data, we used the estimates of life expectancy data calculated by the authors instead of maximum lifespan for species, following what the authors did in their paper. Vincze et al. ^4^ also had only one category for neoplastic conditions represented by any neoplasm that was diagnosed as cause of death while Boddy et al. ^3^ separated total neoplasms and cancer as in our work. As such, the data of Vincze et al. ^4^ represent cancer mortality prevalence, while the data of Boddy et al. ^3^ were analyzed for neoplasia, cancer, and malignancy transformation rate. Note that while Boddy et al. ^3^ did not use their data to calculate the malignancy transformation rate, their data could be used to calculate all the categories used in this paper.

## ACKNOWLEDGMENTS

This project was supported by NSF IOS joint collaborative awards (No. 2028458 to VJL and No. 2028459 to SG and YC). We are grateful to Harald Schmidt for helping us to obtain the data at the Rotterdam Zoo, to Isabella Capellini for providing the avian phylogeny used in our study, and to Beth Reinke for allowing us access to reptile and amphibian life history trait data. We are also grateful to Robert Ossiboff for our discussion on veterinary medicine and pathology reports on different vertebrate groups. Finally, we are thankful to the comments received by two anonymous reviewers, which strongly improved the clarity and potential impact of our paper.

## AUTHOR CONTRIBUTIONS

Conceptualization: YC, SG; Investigation: SB, LP, WA, NM, J-PA, SD, SMcC, PW, SB, LRD, LGRB - vanS, DF, SG, YC; Data curation: SB, SG, YC; Formal Analysis: SB, JC, SG, YC; Writing – Original Draft: SB, SG, YC; Writing – Review and Editing: SMcC, PW, LGRB - vanS, DF, VJL, JC.

## DATA AVAILABILITY

All data and R codes used for this paper will be available on Dryad after manuscript acceptance.

