## Supplementary Materials for "Unraveling the relationship between cancer and life history traits in vertebrates"

**Supplementary Materials Table 1.** Number of species/necropsies per tetrapod group collected from each zoo and squamate snake data from, Duke et al. 2022 *before* secondary curation (see Methods for additional explanation). Since species representation overlaps amongst the zoos, the total species number does not reflect the sum of the species count per zoo. Dashed lines indicate that data for that specific group were not collected (or available) from that zoo. The full dataset can be found on Dryad (*after manuscript acceptance*)

| **Source** | **Amphibians** | **Birds** | **Mammals** | **Crocodilians** | **Turtles** | **Squamates** |
|---|---|---|---|---|---|---|
| Allwetterzoo, Germany | 9/24 | 156/930 | 91/726 | 2/2 | 18/45 | 34/170 |
| Birmingham Zoo, USA | – | – | – | 1/4 | 19/29 | 55/106 |
| Hellenthal Zoo | – | 65/163 | 27/196 | – | – | – |
| Rotterdam Zoo, The Netherlands | 26/162 | 290/3148 | 161/2082 | 6/14 | 36/149 | 112/503 |
| Der Grüner Zoo, Wuppertal, Germany | 14/72 | 190/748 | 67/629 | 1/5 | 11/19 | 34/80 |
| ZOOM Erlebniswelt Gelsenkirchen, Germany | – | 38/152 | 65/387 | – | 4/14 | 10/15 |
| Zürich Zoologisher Garten, Switzerland | 23/137 | 48/448 | – | – | 9/26 | 44/256 |
| Duke et al. 2022 | – | – | – | – | – | 1771 |
| **Total Reports** | 50/395 | 512/5581 | 258/4020 | 8/25 | 60/282 | 172/2901 |

**Supplementary Materials Table 2.** Shannon-Wiener Diversity Index and the Shannon Equitability Index were used to assess species diversity and the evenness of species distribution in our dataset (Shannon, 1948; Tuomisto, 2012). These indices were calculated on the datasets used to run the *gee* analyses. The Shannon-Wiener Diversity Index represents the proportion of species that make up the population of each group and thus the average amount of diversity for the group. The Shannon-Wiener Diversity Index (H) is calculated as follows:

$$H' = - \sum_{i=1}^{S} p_i \ln p_i$$

Where $p_i$ is the relative abundance of species i, S is the total number of species present in the group and ln is the natural log. The evenness was then calculated using the Shannon Equitability Index: $E_H$ = H/ln(S). The Shannon Equitability Index is a measure of how even individuals (i.e., necropsies) are distributed amongst the species of each group. A value of 1 means all species are equally represented by the same number of necropsies, while a value closer to 0 means that one or a small number of species are overrepresented.

The Shannon-Wiener Diversity Index and the Shannon Equitability Index for each studied group are indicated here below. Based on the Shannon-Wiener Diversity Index, birds and mammals had the highest sampled species diversity, but they also had the most necropsies, while crocodilians were underrepresented both in terms of number of species and number of necropsies compared to other tetrapod groups. Even though the total number of species and individuals for each group was different, the Shannon Equitability Index indicated that the allocation of necropsies among species was similar within each of the groups (0.8-0.9).

| | **Amphibians** | **Birds** | **Crocodilians** | **Mammals** | **Squamates** | **Squamates*** | **Turtles** |
|---|---|---|---|---|---|---|---|
| **Diversity Index** | 3.33 | 4.92 | 1.73 | 4.75 | 4.23 | 4.17 | 3.57 |
| **Equitability Index** | 0.866 | 0.861 | 0.888 | 0.879 | 0.876 | 0.839 | 0.907 |

**Supplementary Materials Table 3.** Species listed whose prevalence is significantly ($p = <0.05$) different from the mean for its group. Analysis performed with *prop.test* (R package {stats}). Numbers in parenthesis are the actual numbers of incidence/necropsy number or cancer/neoplasia in the case of malignancy. NA in malignancy denotes there were no significant outliers for that group. Turtles and crocodilians did not have significant outliers from the mean and were not included in the table.

| **Group** | **Neoplasia** | **Cancer** | **Malignancy** |
|---|---|---|---|
| **Amphibians** | *Ambystoma mexicanum* (Axolotl)(3/11) 27% | *Ambystoma mexicanum* (Axolotl)(3/11) 27% | NA |
| | *Leptodactylus fallax* (Mountain Chicken Frog)(3/16) 19% | *Leptodactylus fallax* (Mountain Chicken Frog)(3/16) 19% | |
| | *Pipa pipa* (Surinam toad) (4/6) 67% | | |
| **Birds** | *Bubo bubo* (Eurasian eagle owl) (2/10) 20% | *Cairina moschata* (Muscovy duck)(1/3) 33% | NA |
| | *Cacatua leadbeateri* (Mitchell's cockatoo)(1/2)50% | *Coturnix delegorguei* (Harlequin quail)(2/15)13% | |
| | *Coturnix delegorguei* (Harlequin quail)(2/15)13% | *Dendrocygna viduata* (White-faced whistling duck)(4/48) 8% | |
| | *Dendrocygna viduata* (White-faced whistling duck)(4/48) 8% | *Falco peregrinus* (Peregrine falcon)(2/8)25% | |
| | *Falco peregrinus* (Peregrine falcon)(2/8)25% | *Forpus coelestis* (Pacific parrotlet)(1/2) 50% | |
| | *Forpus coelestis* (Pacific parrotlet)(1/2) 50% | *Furnarius leucopus* (Pale-legged hornero)(1/2) 50% | |
| | *Furnarius leucopus* (Pale-legged hornero)(1/2) 50% | *Gallus gallus* (Domestic chicken)(5/157) 4% | |
| | *Gallus gallus* (Domestic chicken)(7/157)5.6% | *Geococcyx californianus* (Roadrunner)(1/4) 25% | |
| | *Geococcyx californianus* (Roadrunner)(2/4) 50% | *Grus virgo* (Demoiselle crane)(2/9) 22% | |
| | *Grus virgo* (Demoiselle crane)(3/9) 33% | *Jacana jacana* (Wattled jacana)(2/6) 33% | |
| | *Jacana jacana* (Wattled jacana)(2/6) 33% | *Panurus biarmicus* (Bearded tit babbler)(1/4) 25% | |

| | | | |
|---|---|---|---|
| | *Philemon citreogularis* (Little friarbird)(1/1) 100% | *Philemon citreogularis* (Little friarbird)(1/1) 100% | |
| | *Ptilinopus pulchellus* (Beautiful fruit dove)(1/1) 100% | *Rhea americana* (Greater rhea)(1/4) 25% | |
| | *Pycnonotus cafer* (Red-vented bulbul)(2/2) 100% | *Rupicola peruvianus* (Andean cock-of-the-rock)(2/8) 25% | |
| | *Rupicola peruvianus* (Andean cock-of-the-rock)(2/8) 25% | *Tadorna ferruginea* (Ruddy schelduck)(1/2) 50% | |
| | *Tadorna ferruginea* (Ruddy schelduck)(1/2) 50% | *Tauraco leucotis* (White-cheeked turaco)(1/1) 100% | |
| | *Tauraco leucotis* (White-cheeked turaco)(1/1) 100% | *Tragopan satyra* (Satyr tragopan)(1/1) 100% | |
| | *Tragopan satyra* (Satyr tragopan)(1/1) 100% | | |
| **Mammals** | *Callimico goeldii* (Geoldi's marmoset)(4/8) 50% | *Callimico goeldii* (Geoldi's marmoset)(2/8) 25% | *Catopuma temminckii* (Asian gold cat)(10/10) 100% |
| | *Capra hircus* (Domestic goat)(0/91) 0% | *Catopuma temminckii* (Asian gold cat)(10/25) 40% | *Cricetus cricetus* (Common hamster) (8/25) 32% |
| | *Catopuma temminckii* (Asian gold cat)(10/25) 40% | *Dasyuroides byrnei* (Kowari)(2/2) 100% | |
| | *Cercopithecus cephus* (Moustached guenon)(2/2) 100% | *Lynx lynx* (Eurasian lynx)(2/5) 40% | |
| | *Chrysocyon brachyurus* (Maned wolf)(2/5) 40% | *Martes flavigula* (Yellow-throated marten)(2/8) 25% | |
| | *Cricetus cricetus* (Common hamster) (25/114) 22% | *Microcebus murinus* (Grey mouse lemur)(3/12) 25% | |
| | *Dasyuroides byrnei* (Kowari)(2/2) 100% | *Panthera leo* (Lion)(6/27) 22% | |
| | *Dolichotis patagonum* (Patagonian mara)(0/103) 0% | *Panthera pardus* (Leopard)(4/14) 29% | |

| | | | |
|---|---|---|---|
| | *Lynx lynx* (Eurasian lynx)(2/5) 40% | *Prionailurus viverrinus* (Fishing cat)(3/12) 25% | |
| | *Mephitis mephitis* (Striped skunk)(3/7) 43% | *Procyon lotor* (Raccoon)(8/30) 27% | |
| | *Microcebus murinus* (Grey mouse lemur)(5/12) 42% | *Ursus arctos* (Brown Bear)(2/5) 40% | |
| | *Panthera leo* (Lion)(8/27) 30% | *Ursus maritimus* (Polar bear)(2/6) 33% | |
| | *Panthera pardus* (Leopard)(5/14) 36% | | |
| | *Pecari tajacu* (Collared peccary)(0/89) 0% | | |
| | *Phacochoerus africanus* (Common warthog)(4/10) 40% | | |
| | *Procyon lotor* (Raccoon)(9/30) 30% | | |
| | *Rattus norvegicus* (Brown rat)(4/10) 40% | | |
| | *Ursus arctos* (Brown Bear)(2/5) 40% | | |
| **Squamates** | *Epicrates fordi* (Haiti ground boa)(4/5) 80% | *Crotalus cerastes (Sonoran desert sidewinder)*(2/5) 40% | NA |
| | *Orthriophis taeniurus* (Beauty snake)(4/15) 26% | *Epicrates fordi* (Haiti ground boa)(3/5) 60% | |
| | *Pantherophis guttatus* (Corn snake)(5/35) 14% | *Orthriophis taeniurus* (Beauty snake)(4/15) 26% | |
| | *Varanus dumerilii* (Dumeril's monitor)(2/4) 50% | *Varanus dumerilii* (Dumeril's monitor)(2/4) 50% | |
| **Squamates*** | Crotalus adamanteus (Eastern diamondback rattlesnake)(6/21) 29% | Crotalus adamanteus (Eastern diamondback rattlesnake)(6/21) 29% | *Cyclura cornuta* (Rhinoceros iguana)(0/2) 0% |
| | *Crotalus cerastes (Sonoran desert sidewinder)*(3/10) 30% | *Crotalus cerastes (Sonoran desert sidewinder)*(3/10) 30% | *Pogona vitticeps* (Inland bearded dragon)(0/2) 0% |
| | *Crotalus horridus* (Timber rattlesnake)(5/24) 21% | *Epicrates fordi* (Haiti ground boa)(3/5) 60% | *Thamnophis exsul* (Exiled Gartersnake)(0/2) 0% |

| | | | |
|---|---|---|---|
| | *Epicrates fordi* (Haiti ground boa)(4/5) 80% | *Lampropeltis getula* (Kingsnake)(5/28) 18% | *Tiliqua rugosa* (Shingleback skink)(0/2) 0% |
| | *Nerodia sipedon* (Northern water snake)(4/14) 29% | *Nerodia sipedon* (Northern water snake)(4/14) 29% | |
| | *Orthriophis taeniurus* (Beauty snake)(5/24) 21% | *Orthriophis taeniurus* (Beauty snake)(5/24) 21% | |
| | *Varanus dumerilii* (Dumeril's monitor)(2/4) 50% | *Varanus dumerilii* (Dumeril's monitor)(2/4) 50% | |

***** Includes data from Duke et. al (2022)

**Supplementary Materials Table 4.** Dataset numbers only for species with 10 or more and 20 or more necropsies. "Total neoplasia #" includes all benign and malignant tumor counts whereas "Total cancer #" include only those tumors that were diagnosed as a malignancy by a veterinary pathologist and confirmed by histology. Malignancy prevalence is derived from the cancer count out of neoplasia count. Amphibians, crocodilians, and turtles were not included in 10+ and 20+ analysis due to the total number of species being less than 20, which we considered as the minimum number for the number of species to obtain reliable results.

a) 10+ necropsies

| | **Birds** | **Mammals** | **Squamates** | **Squamates*** |
|---|---|---|---|---|
| **# Species with >9 necropsies** | 124 | 80 | 28 | 64 |
| **# of necropsies** | 3395 | 2255 | 575 | 2136 |
| **Total neoplasia #** | 49 | 133 | 45 | 159 |
| **Cancer count** | 31 | 81 | 29 | 133 |
| **Prevalence neoplasia** | 0.014 | 0.059 | 0.078 | 0.074 |
| **Prevalence cancer** | 0.009 | 0.036 | 0.05 | 0.062 |
| **Prevalence malignancy** | 0.633 | 0.61 | 0.644 | 0.84 |

***** Includes data from Duke et. al (2022)

b) 20+ necropsies

| | **Birds** | **Mammals** | **Squamates*** |
|---|---|---|---|
| **# Species with >19 necropsies** | 55 | 39 | 32 |
| **# of necropsies** | 2735 | 1658 | 1488 |
| **Total neoplasia #** | 44 | 89 | 101 |
| **Cancer count** | 30 | 55 | 88 |
| **Prevalence neoplasia** | 0.016 | 0.054 | 0.07 |
| **Prevalence cancer** | 0.01 | 0.03 | 0.06 |
| **Prevalence malignancy** | 0.7 | 0.6 | 0.9 |

***** Includes data from Duke et. al (2022)

**Supplementary Materials Table 5.** Pairwise comparison results for (a) neoplasia, (b) cancer, and (c) malignancy prevalence proportions, run on species with 10 or more necropsies using *prop.test and p.adjust* with the Benjamini & Hochberg ("BH") method (R package {stats}) (R package {stats}). Numbers in parentheses are proportion numbers: neoplasia and cancer proportions derived from total necropsy number while malignancy is the proportion of cancer from neoplasia. Significant values are in bold. Malignancy could not be calculated for crocodile since there were no instances of cancer in this group in species with 10 or more necropsies. Squamates* includes data from Duke et. al (2022).

a) GROUP NEOPLASIA PREVALENCE

| | Amphibians (7/266) | Birds (56/3872) | Mammals (133/2255) | Squamates (45/575) | Squamates* (159/2136) | Turtles (1/67) |
|---|---|---|---|---|---|---|
| Turtles (1/67) | 0.98 | 1 | 0.26 | **8.25e-16** | 0.16 | 1 |
| Squamates* (159/2136) | **0.015** | **8.25e-16** | 0.086 | 0.95 | 1 | |
| Squamates (45/575) | **0.015** | **8.25e-16** | 0.16 | 1 | | |
| Mammals (133/2255) | 0.08 | **8.25e-16** | 1 | | | |
| Birds (56/3872) | 0.26 | 1 | | | | |

b) GROUP CANCER PREVALENCE

| | Amphibians (7/266) | Birds (35/3872) | Mammals (81/2255) | Squamates (29/575) | Squamates*( 133/2136) | Turtles (0/67) |
|---|---|---|---|---|---|---|
| Turtles (0/67) | **1.73E-14** | 0.9 | 0.27 | 0.19 | 0.12 | 1 |
| Squamates* (133/2136) | 0.06 | **3.3E-15** | **0.0002** | 0.4 | 1 | |
| Squamates (29/575) | 0.21 | **1.6E-13** | 0.21 | 1 | | |
| Mammals (81/2255) | 0.57 | **7.5E-13** | 1 | | | |
| Birds (35/3872) | **0.041** | 1 | | | | |

c) GROUP MALIGNANCY PREVALENCE

| | Amphibians (7/7) | Birds (35/56) | Mammals (81/133) | Squamates (29/45) | Squamates* (133/159) | Turtles (0/1) |
|---|---|---|---|---|---|---|
| Turtles (0/1) | 0.48 | **0.96** | **0.96** | **0.96** | 0.7 | 1 |
| Squamates* (133/159) | 0.88 | **0.014** | **0.0003** | **0.05** | 1 | |
| Squamates (29/45) | 0.36 | 1 | 0.96 | 1 | | |
| Mammals (81/133) | 0.35 | 1 | 1 | | | |
| Birds (35/56) | 0.36 | 1 | | | | |

**Supplementary Materials Table 6.** Pairwise comparison results for (d) neoplasia, cancer (e), and (f) malignancy prevalence proportions, run on species with 20 or more necropsies using *prop.test and p.adjust* with the Benjamini & Hochberg ("BH") method (R package {stats}) (R package {stats}). Numbers in parentheses are proportion numbers: neoplasia and cancer proportions derived from total necropsy number while malignancy is the proportion of cancer from neoplasia. Significant values are in bold. Turtles and crocodiles were not included in this table as they did not have any species with 20 or more necropsy reports. Squamates* includes data from Duke et. al (2022)

d) GROUP NEOPLASIA PREVALENCE

| | Amphibians (1/167) | Birds (45/2998) | Mammals (93/1732) | Squamates (22/327) | Squamates* (122/1780) |
|---|---|---|---|---|---|
| Squamates* (122/1780) | **0.007** | **2.2e-15** | 0.11 | 1 | 1 |
| Squamates (22/327) | **0.009** | **2.14e-9** | 0.5 | 1 | |
| Mammals (93/1732) | **0.02** | **2.61e-13** | 1 | | |
| Birds (45/2998) | 0.6 | 1 | | | |

e) GROUP CANCER PREVALENCE

| | Amphibians (1/167) | Birds (31/2998) | Mammals (59/1732) | Squamates (15/327) | Squamates*(107/1780) |
|---|---|---|---|---|---|
| Squamates* (107/1780) | **0.012** | **2.2E-15** | **0.0009** | 0.42 | 1 |
| Squamates (15/327) | 0.06 | **2.2E-6** | 0.42 | 1 | |
| Mammals (59/1732) | 0.12 | **8.4E-8** | 1 | | |
| Birds (31/2998) | 0.88 | 1 | | | |

f) GROUP MALIGNANCY PREVALENCE

| | Amphibians (1/1) | Birds (31/45) | Mammals (59/93) | Squamates (15/22) | Squamates* (107/122) |
|---|---|---|---|---|---|
| Squamates* (107/122) | 1 | **0.044** | **0.0005** | 0.14 | 1 |

| | | | | |
|---|---|---|---|---|
| Squamates (15/22) | 1 | 1 | 1 | 1 |
| Mammals (59/93) | 1 | 1 | 1 | |
| Birds (31/45) | 1 | 1 | | |

**Supplementary Materials Table 7.** Neoplasia, cancer, and malignancy prevalence breakdown by family or order within each tetrapod group. Mammals and birds were broken down by order, while all the others by family. Malignancy prevalence of 50% or higher and the highest neoplasia and cancer prevalences in each group are highlighted in bold. Significance of differences in the proportion of prevalence among families (for amphibians, squamate) or orders (for birds and mammals) within each group were tested in pairwise comparison using *par.test* in R. Results of pairwise comparisons can be found on Dryad *after manuscript acceptance*.

**Summary of Results P-values derived from pairwise comparison by family (order for mammals and birds) within each group:**
In amphibians, Pipidae (30%) and Ambystomatidae (27%) had the highest neoplasia prevalence values, which were significantly ($p<0.05$) different from other major amphibian families including Bufonidae, Dendrobatidae, Hylidae, Microhylidae, and Salamandridae. Ten amphibian families had 0% neoplasia prevalence, and Bufonidae had the lowest non-zero prevalence (3%).

In turtles, the only neoplasm was found in *Chelus fimbriata* (Chelidae). In crocodilians, there were only two incidences of neoplasia, one each in Alligatoridae and Crocodylidae, with malignancy transformation only occurring in *Osteolaemus tetraspis* (Crocodylidae).

In squamates, the highest neoplasia prevalence was found in the Helodermatidae (25%) and was significantly ($p<0.05$) different from several other squamate families. Malignancy transformation rate was high in snakes, especially in the Colubridae (70%) and Viperidae (100%).

Cancer was widespread across mammals, with 10 out of 17 orders represented in our work experiencing neoplasia and cancer. The highest prevalence values occurred in carnivores (15% neoplasia and 80% malignancy transformation rate) and primates (11% neoplasia and 48% malignancy transformation rate). In most cases in which neoplasia was detected in mammals, malignancy transformation rate was also high (>30%) with the exception of Chiroptera (4% neoplasia and 0% malignancy transformation) and Proboscidea (25% neoplasia and 0% malignancy transformation).

In birds, although neoplasia was found to occur across several orders with low prevalence, higher neoplasia prevalence was found in Cuculiformes (18.2%), with significant pairwise differences from most other avian orders. When malignancy transformation did occur in birds, the prevalence was extremely high (>50%), except for Pelecaniformes (40%) and Phoenicopteriformes (33%).

**Amphibians**

| Order | Family | # Species in dataset | Total necropsy | Total neoplasias | Total cancer | Neoplasia prevalence | Cancer prevalence | Malignancy prevalence |
|---|---|---|---|---|---|---|---|---|
| **Anura** | Aromobatidae | 1 | 1 | 0 | 0 | 0.000 | 0.000 | 0 |
| | Bombinatoridae | 2 | 22 | 1 | 1 | 0.045 | 0.045 | **1.000** |
| | Bufonidae | 8 | 68 | 2 | 0 | 0.029 | 0.000 | 0.000 |
| | Dendrobatidae | 13 | 101 | 0 | 0 | 0 | 0.000 | 0 |
| | Hemiphractidae | 1 | 9 | 0 | 0 | 0 | 0.000 | 0 |
| | Hylidae | 3 | 36 | 0 | 0 | 0 | 0.000 | 0 |
| | **Leptodactylidae** | 1 | 16 | 3 | 3 | **0.188** | **0.188** | **1.000** |
| | Mantellidae | 4 | 18 | 0 | 0 | 0 | 0.000 | 0 |

| | | | | | | | | |
|---|---|---|---|---|---|---|---|---|
| | Microhylidae | 2 | 35 | 0 | 0 | 0 | 0.000 | 0 |
| | Pelodryadidae | 1 | 7 | 1 | 1 | 0.143 | 0.143 | **1.000** |
| | Pipidae | 2 | 12 | 4 | 1 | 0.3 | 0.083 | 0.250 |
| | Rana | 3 | 12 | 0 | 0 | 0.0 | 0.000 | 0 |
| | Telmatobiidae | 1 | 1 | 0 | 0 | 0 | 0.000 | 0 |
| Anura Total | | 42 | 338 | 11 | 6 | 0.033 | 0.018 | **0.545** |
| **Urodela** | **Ambystomatidae** | 1 | 11 | 3 | 3 | **0.273** | **0.273** | **1.000** |
| | Cryptobranchidae | 1 | 6 | 1 | 1 | 0.167 | 0.167 | **1.000** |
| | Salamandridae | 4 | 27 | 0 | 0 | 0 | 0.000 | 0 |
| | Sirenidae | 1 | 1 | 0 | 0 | 0 | 0.000 | 0 |
| Urodela Total | | 7 | 45 | 4 | 4 | 0.089 | 0.089 | **1.000** |
| Grand Total | | 49 | 383 | 15 | 10 | 0.039 | 0.026 | **0.667** |

### Turtles

| Group | Family | # Species in dataset | Total necropsy | Total neoplasias | Total cancer | Neoplasia prevalence | Cancer prevalence | Malignancy prevalence |
|---|---|---|---|---|---|---|---|---|
| **turtle** | **Chelidae** | 1 | 13 | 1 | 0 | **0.077** | 0.000 | 0 |
| | Cheloniidae | 3 | 3 | 0 | 0 | 0 | 0.000 | 0 |
| | Chelydridae | 2 | 5 | 0 | 0 | 0 | 0.000 | 0 |
| | Emydidae | 8 | 26 | 0 | 0 | 0 | 0.000 | 0 |
| | Geoemydidae | 15 | 32 | 0 | 0 | 0 | 0.000 | 0 |
| | Pelomedusidae | 2 | 3 | 0 | 0 | 0 | 0.000 | 0 |
| | Podocnemididae | 2 | 4 | 0 | 0 | 0 | 0.000 | 0 |
| | Testudinidae | 23 | 120 | 0 | 0 | 0 | 0.000 | 0 |
| | Trionychidae | 1 | 1 | 0 | 0 | 0 | 0.000 | 0 |
| Grand Total | | 57 | 158 | 1 | 0 | 0.006 | 0.000 | 0 |

### Squamates

| Group | Family | # Species in dataset | Total necropsy | Total neoplasias | Total cancer | Neoplasia prevalence | Cancer prevalence | Malignancy prevalence |
|---|---|---|---|---|---|---|---|---|
| **lizard** | Agamidae | 19 | 98 | 7 | 3 | 0.071 | 0.031 | 0.429 |
| | Chamaeleonidae | 8 | 62 | 1 | 1 | 0.016 | 0.016 | **1.000** |
| | Cordylidae | 5 | 15 | 1 | 1 | 0.0667 | 0.067 | **1.000** |
| | Corytophanidae | 2 | 26 | 0 | 0 | 0 | 0.000 | 0.000 |
| | Crotaphytidae | 1 | 7 | 0 | 0 | 0 | 0.000 | 0.000 |
| | Dactyloidae | 7 | 36 | 0 | 0 | 0 | 0.000 | 0.000 |
| | **Diplodactylidae** | 2 | 7 | 1 | 1 | 0.143 | **0.143** | **1.000** |
| | Eublepharidae | 2 | 10 | 1 | 0 | 0.1 | 0.000 | 0.000 |

| Group | Family | # Species in dataset | Total necropsy | Total neoplasias | Total cancer | Neoplasia prevalence | Cancer prevalence | Malignancy prevalence |
|---|---|---|---|---|---|---|---|---|
| | Gekkonidae | 11 | 80 | 1 | 0 | 0.013 | 0.000 | 0.000 |
| | Gerrhosauridae | 2 | 15 | 0 | 0 | 0 | 0.000 | 0.000 |
| | **Helodermatidae** | 2 | 12 | 3 | 1 | **0.250** | 0.083 | 0.333 |
| | Iguanidae | 10 | 68 | 6 | 4 | 0.088 | 0.059 | **0.667** |
| | Lacertidae | 4 | 17 | 1 | 1 | 0.059 | 0.059 | **1.000** |
| | Opluridae | 3 | 14 | 1 | 1 | 0.0714 | 0.071 | 0.000 |
| | Phrynosomatidae | 7 | 21 | 0 | 0 | 0 | 0.000 | 0.000 |
| | Polychrotidae | 1 | 13 | 0 | 0 | 0 | 0.000 | 0.000 |
| | Scincidae | 12 | 64 | 5 | 2 | 0.078 | 0.031 | 0.400 |
| | Shinisauria | 1 | 16 | 2 | 1 | 0.125 | 0.063 | **0.500** |
| | Sphaerodactylidae | 1 | 5 | 1 | 0 | 0.2 | 0.000 | 0.000 |
| | Teiidae | 3 | 8 | 0 | 0 | 0 | 0.000 | 0.000 |
| | Varanidae | 14 | 68 | 6 | 6 | 0.088 | 0.088 | **1.000** |
| lizard Total | | 117 | 662 | 37 | 22 | 0.056 | 0.033 | **0.595** |
| **snake** | Boidae | 12 | 78 | 9 | 6 | 0.115 | 0.077 | **0.667** |
| | Colubridae | 22 | 221 | 26 | 19 | 0.118 | 0.086 | **0.731** |
| | **Elapidae** | 3 | 3 | 1 | 1 | **0.333** | **0.333** | **1.000** |
| | Lamprophiidae | 1 | 1 | 0 | 0 | 0 | 0.000 | 0.000 |
| | Pythonidae | 10 | 59 | 7 | 5 | 0.119 | 0.085 | **0.714** |
| | Viperidae | 14 | 29 | 4 | 4 | 0.138 | 0.138 | **1.000** |
| snake Total | | 62 | 391 | 47 | 35 | 0.120 | 0.090 | **0.745** |
| Grand Total | | 179 | 1053 | 84 | 57 | 0.080 | 0.054 | **0.679** |

### Squamates*

| Group | Family | # Species in dataset | Total necropsy | Total neoplasias | Total cancer | Neoplasia prevalence | Cancer prevalence | Malignancy prevalence |
|---|---|---|---|---|---|---|---|---|
| **lizard** | Agamidae | 19 | 98 | 7 | 3 | 0.071 | 0.031 | 0.429 |
| | Chamaeleonidae | 8 | 62 | 1 | 1 | 0.016 | 0.016 | **1.000** |
| | Cordylidae | 5 | 15 | 1 | 1 | 0.0667 | 0.067 | **1.000** |
| | Corytophanidae | 2 | 26 | 0 | 0 | 0 | 0.000 | 0.000 |
| | Crotaphytidae | 1 | 7 | 0 | 0 | 0 | 0.000 | 0.000 |
| | Dactyloidae | 7 | 36 | 0 | 0 | 0 | 0.000 | 0.000 |
| | Diplodactylidae | 2 | 7 | 1 | 1 | 0.143 | 0.143 | **1.000** |
| | Eublepharidae | 2 | 10 | 1 | 0 | 0.1 | 0.000 | 0.000 |
| | Gekkonidae | 11 | 80 | 1 | 0 | 0.013 | 0.000 | 0.000 |
| | Gerrhosauridae | 2 | 15 | 0 | 0 | 0 | 0.000 | 0.000 |

| | | | | | | | | |
|---|---|---|---|---|---|---|---|---|
| | Helodermatidae | 2 | 12 | 3 | 1 | 0.250 | 0.083 | 0.333 |
| | Iguanidae | 10 | 68 | 6 | 4 | 0.088 | 0.059 | **0.667** |
| | Lacertidae | 4 | 17 | 1 | 1 | 0.059 | 0.059 | **1.000** |
| | Opluridae | 3 | 14 | 1 | 1 | 0.071 | 0.071 | 0.000 |
| | Phrynosomatidae | 7 | 21 | 0 | 0 | 0 | 0.000 | 0.000 |
| | Polychrotidae | 1 | 13 | 0 | 0 | 0 | 0.000 | 0.000 |
| | Scincidae | 12 | 64 | 5 | 2 | 0.078 | 0.031 | 0.400 |
| | Shinisauria | 1 | 16 | 2 | 1 | 0.125 | 0.063 | **0.500** |
| | Sphaerodactylidae | 1 | 5 | 1 | 0 | 0.2 | 0.000 | 0.000 |
| | Teiidae | 3 | 8 | 0 | 0 | 0 | 0.000 | 0.000 |
| | Varanidae | 14 | 68 | 6 | 6 | 0.088 | 0.088 | **1.000** |
| lizard Total | | 117 | 662 | 37 | 22 | 0.056 | 0.033 | **0.595** |
| **snake** | **Boidae** | 16 | 193 | 18 | 13 | **0.093** | 0.067 | **0.722** |
| | Colubridae | 35 | 1027 | 86 | 76 | 0.084 | 0.074 | **0.884** |
| | Elapidae | 6 | 128 | 7 | 7 | 0.055 | 0.055 | **1.000** |
| | Lamprophiidae | 1 | 1 | 0 | 0 | 0.000 | 0.000 | 0 |
| | Pythonidae | 12 | 203 | 15 | 13 | 0.074 | 0.064 | **0.867** |
| | **Viperidae** | 22 | 394 | 35 | 33 | 0.089 | **0.084** | **0.943** |
| snake Total | | 92 | 1946 | 161 | 142 | 0.083 | 0.073 | **0.882** |
| Grand Total | | 209 | 2608 | 198 | 164 | 0.076 | 0.063 | **0.828** |

### Mammals

| Order | # Species in dataset | Total necropsy | Total neoplasias | Total cancer | Neoplasia prevalence | Cancer prevalence | Malignancy prevalence |
|---|---|---|---|---|---|---|---|
| Afrosoricida | 2 | 5 | 0 | 0 | 0.000 | 0.000 | 0.000 |
| Artiodactyla | 67 | 1021 | 34 | 16 | 0.033 | 0.016 | 0.471 |
| **Carnivora** | 53 | 532 | 78 | 63 | **0.147** | 0.118 | **0.808** |
| Chiroptera | 4 | 26 | 1 | 0 | 0.038 | 0.000 | 0.000 |
| **Dasyuromorphia** | 1 | 2 | 2 | 2 | **1** | **1.000** | **1.000** |
| Diprotodontia | 12 | 136 | 4 | 1 | 0.029 | 0.007 | 0.250 |
| Eulipotyphla | 1 | 1 | 0 | 0 | 0 | 0.000 | 0.000 |
| Hyracoidae | 1 | 48 | 1 | 1 | 0.021 | 0.021 | **1.000** |
| Lagomorpha | 2 | 61 | 0 | 0 | 0 | 0.000 | 0.000 |
| Macroscelidea | 3 | 52 | 0 | 0 | 0 | 0.000 | 0.000 |
| Monotremata | 1 | 2 | 0 | 0 | 0 | 0.000 | 0.000 |
| Perissodactyla | 8 | 50 | 3 | 2 | 0.060 | 0.040 | **0.667** |
| Pholidota | 1 | 2 | 0 | 0 | 0.000 | 0.000 | 0.000 |

| | | | | | | | |
|---|---|---|---|---|---|---|---|
| **Primate** | 40 | 261 | 29 | 14 | **0.111** | 0.054 | 0.483 |
| **Proboscidea** | 1 | 4 | 1 | 0 | **0.250** | 0.000 | 0.000 |
| Rodentia | 31 | 636 | 38 | 12 | 0.060 | 0.019 | 0.316 |
| Scandentia | 2 | 2 | 0 | 0 | 0.000 | 0.000 | 0.000 |
| Grand Total | 230 | 2841 | 191 | 111 | 0.067 | 0.039 | **0.581** |

**Birds**

| Order | # Species in dataset | Total necropsy | Total neoplasias | Total cancer | Neoplasia prevalence | Cancer Prevalence | Malignancy prevalence |
|---|---|---|---|---|---|---|---|
| Accipitriformes | 32 | 121 | 2 | 1 | 0.017 | 0.008 | **0.5** |
| Anseriformes | 86 | 1217 | 22 | 14 | 0.018 | 0.012 | **0.636** |
| Apodiformes | 4 | 9 | 0 | 0 | 0.000 | 0.000 | 0 |
| Bucerotiformes | 12 | 33 | 2 | 0 | 0.061 | 0.000 | 0 |
| Cariamiformes | 1 | 11 | 0 | 0 | 0.000 | 0.000 | 0 |
| Casuariiformes | 1 | 4 | 0 | 0 | 0 | 0.000 | 0 |
| Charadriiformes | 23 | 395 | 6 | 6 | 0.015 | 0.015 | **1.000** |
| Ciconiiformes | 6 | 40 | 0 | 0 | 0 | 0.000 | 0 |
| Coliiformes | 1 | 45 | 0 | 0 | 0 | 0.000 | 0 |
| Columbiformes | 25 | 177 | 2 | 0 | 0.011 | 0.000 | 0 |
| Coraciiformes | 13 | 78 | 0 | 0 | 0 | 0.000 | 0 |
| **Cuculiformes** | 5 | 11 | 2 | 1 | **0.182** | **0.091** | **0.5** |
| Eurypygiformes | 1 | 3 | 0 | 0 | 0 | 0.000 | 0 |
| Falconiformes | 8 | 38 | 2 | 2 | 0.053 | 0.053 | **1.000** |
| Galliformes | 40 | 435 | 16 | 10 | 0.037 | 0.023 | **0.625** |
| Gruiformes | 11 | 70 | 3 | 2 | 0.043 | 0.029 | **0.667** |
| Musophagiformes | 11 | 75 | 2 | 1 | 0.027 | 0.013 | **0.5** |
| Passeriformes | 122 | 1089 | 11 | 7 | 0.010 | 0.006 | **0.636** |
| Pelecaniformes | 18 | 313 | 5 | 2 | 0.016 | 0.006 | 0.4 |
| Phoenicopteriformes | 4 | 86 | 3 | 1 | 0.035 | 0.012 | 0.333 |
| Piciformes | 12 | 105 | 1 | 1 | 0.010 | 0.010 | 0 |
| Podargiformes | 1 | 6 | 0 | 0 | 0 | 0.000 | 0 |
| Podicipediformes | 2 | 6 | 0 | 0 | 0 | 0.000 | 0 |
| Procellariiformes | 1 | 1 | 0 | 0 | 0 | 0.000 | 0 |
| Psittaciformes | 39 | 202 | 4 | 3 | 0.020 | 0.015 | **0.75** |
| Pterocliformes | 1 | 2 | 0 | 0 | 0.000 | 0.000 | 0 |
| Rheiformes | 2 | 32 | 1 | 1 | 0.031 | 0.031 | **1** |
| Sphenisciformes | 4 | 247 | 3 | 3 | 0.012 | 0.012 | **1** |

| Order | # Species in dataset | Total necropsy | Total neoplasias | Total cancer | Neoplasia prevalence | Cancer Prevalence | Malignancy prevalence |
|---|---|---|---|---|---|---|---|
| Strigiformes | 14 | 192 | 7 | 4 | 0.036 | 0.021 | **0.571** |
| Struthioniformes | 1 | 24 | 0 | 0 | 0 | 0.000 | 0 |
| Suliformes | 1 | 22 | 1 | 1 | 0.045 | 0.045 | **1** |
| Tinamiformes | 1 | 33 | 1 | 1 | 0.030 | 0.030 | **1** |
| Trogoniformes | 2 | 3 | 0 | 0 | 0.000 | 0.000 | 0 |
| Grand Total | 505 | 5125 | 96 | 61 | 0.019 | 0.012 | **0.635** |

**Crocodilians**

| Family | # Species in dataset | Total necropsy | Total neoplasias | Total cancer | Neoplasia prevalence | Cancer prevalence | Malignancy prevalence |
|---|---|---|---|---|---|---|---|
| Alligatoridae | 4 | 13 | 1 | 0 | 0.077 | 0.000 | 0 |
| Caimaninae | 1 | 1 | 0 | 0 | 0 | 0.000 | 0 |
| Crocodylidae | 3 | 12 | 1 | 1 | 0.083 | 0.083 | **1** |
| Grand Total | 8 | 26 | 2 | 1 | 0.077 | 0.038 | **0.5** |

**Supplementary Materials Table 8. Influence of body mass or lifespan on cancer, neoplasia, or malignancy prevalence using species with 10 or more and 20 or more necropsies.** Analyses run on species with (a) 10 or more and c) 20 or more necropsies on cancer, neoplasia, and malignancy using either lifespan or body mass as the predicting variable and taking into account group's phylogenetic relationships. Analyses were run using the generalized estimating equations (*gee*; see Materials and Methods for additional information). P-values <0.05 are in bold with slope coefficients in parentheses to show direction effect and jackknife 95% confidence intervals italicized in brackets. Dashed lines represent models that terminate due to non-convergance. Amphibians, crocodiles, and turtles were dropped in these analyses as they did not have enough species with 10 or 20 or more necropsy reports to run models on.

a) 10+ necropsies

| **Vertebrate Group** | **Total Neoplasia** | | | | **Cancer** | | | | **Malignancy** | | | |
|---|---|---|---|---|---|---|---|---|---|---|---|---|
| | **Mass** | | **Lifespan** | | **Mass** | | **Lifespan** | | **Mass** | | **Lifespan** | |
| | **N Species** | **p-value (slope)** ***[jack knife]*** | **N Species** | **p-value (slope)** ***[jack knife]*** | **N Species** | **p-value (slope)** ***[jack knife]*** | **N Species** | **p-value (slope)** ***[jack knife]*** | **N Species** | **p-value (slope)** ***[jack knife]*** | **N Species** | **p-value (slope)** ***[jack knife]*** |
| Birds | 99 | 0.6 (0.04) *[-0.1 , 0.2]* | 86 | 0.9 (-1.1) *[-0.7 , -0.1]* | 99 | — | 86 | **0.004 (-1.1)** ***[-1.3 , -1.1]*** | 30 | 0.3 (0.3) *[0.2 , 0.4]* | 28 | 0.7 (-0.3) *[-0.7 , -0.2]* |
| Mammals | 77 | 0.5 (0.06) *[-0.1 , 0.1]* | 76 | **0.006 (-1.05)** ***[-1.2 , -0.9]*** | 77 | 0.8 (0.03) *[0.02 , 0.04]* | 76 | 0.4 (-0.5) *[-0.5 , -0.2]* | 38 | 0.4 (-0.09) *[-0.1 , -0.04]* | 38 | **0.014 (0.8)** ***[0.8 , 0.9]*** |
| Squamates | 22 | 0.4 (0.08) *[0.02 , 0.3}* | 24 | — | 22 | 0.5 (0.09) *[-0.06 , 0.2]* | 24 | — | 15 | 0.1 (-0.5) *[-0.6, -0.4]* | 15 | 0.7 (-0.5) *[-0.7 , 0.04]* |
| Squamates* | 42 | 0.1 (0.1) *[-0.004 , 0.4]* | 50 | — | 42 | 0.08 (0.2) —- | 50 | — | 35 | 0.6 (0.1) *[0.05 , 0.2]* | 35 | 0.7 (0..3) *[-0.02 , 0.6]* |

b) 20+ necropsies

| Vertebrate Group | **Total Neoplasia** | | | | **Cancer** | | | | **Malignancy** | | | |
|---|---|---|---|---|---|---|---|---|---|---|---|---|
| | **Mass** | | **Lifespan** | | **Mass** | | **Lifespan** | | **Mass** | | **Lifespan** | |
| | **N Species** | **p-value (slope) *[jack knife]*** | **N Species** | **p-value (slope) *[jack knife]*** | **N Species** | **p-value (slope) *[jack knife]*** | **N Species** | **p-value (slope) *[jack knife]*** | **N Species** | **p-value (slope) *[jack knife]*** | **N Species** | **p-value (slope) *[jack knife]*** |
| Birds | 54 | — | 47 | — | 54 | — | 47 | — | 25 | 0.35 (0.3) *[0.2 , 0.4]* | 23 | 0.7 (-0.3) *[-0.8 , 0.2]* |
| Mammals | 39 | 0.6 (0.07) *[0.02 , 0.1]* | 39 | — | 39 | — | 39 | — | 19 | 0.4 (-0.1) *[-0.5 , 0.02]* | 19 | **0.03 (0.9) *[-1.7 , 1.2]*** |
| Squamates* | 24 | 0.8 (0.02) *[-0.09 , 0.1]* | 24 | **0.003 (2.3) *[1.4 , 2.8]*** | 24 | 0.9 (-0.008)*[-0.09 , 0.1]* | 24 | **0.001 (2.9) *[2.6 , 3.0]*** | 22 | 0.9 (-0.0008) *[-0.2 , 0.1]* | 30 | 0.6 (0.4) *[0.2 , 0.5]* |

**Supplementary Materials Table 9.** Re-analysis using *gee* of previously published datasets. Data collected from Boddy et al (2020) and Vincze et al (2022) were analyzed separately using the same *gee* model as used on our own data. Boddy et al. (2022) did not have data on malignancy calculated as in this study (#cancer/#neoplasia), while data from Vincze et al. (2022) represent cancer mortality prevalence.
"**-**" symbol indicates that the analysis did not converge and thus could not produce a result.

| **Dataset** | **N Species** | **Total Neoplasia** | | **Cancer** | | **Malignancy** | |
|---|---|---|---|---|---|---|---|
| | | **Mass p-value (slope) [*jackknife*]** | **Lifespan p-value (slope) [*jackknife*]** | **Mass p-value (slope) [*jackknife*]** | **Lifespan p-value (slope) [*jackknife*]** | **Mass p-value (slope) [*jackknife*]** | **Lifespan p-value (slope) [*jackknife*]** |
| Boddy et al | 36 | 0.4 (0.1) [*-0.01* , *0.1*] | 0.7 (-0.3) [*-0.5* , *0.09*] | 0.8 (0.03) [*-0.02* , *0.07*] | 0.4 (-0.4) [*-0.8* , *0.01*] | — | 0.3 (-0.6) [*-0.9* , *-0.3*] |
| | | **Cancer Mortality** | | | | | |
| | | **Mass p-value (slope) [*jackknife*]** | **Lifespan p-value (slope) [*jackknife*]** | | | | |
| Vincze et al | 189 | **9e-6 (0.2) [*0.18* , *0.22*]** | **1e-4 (1.08) *[0.98 , 1.1]*** | NA | NA | NA | NA |

| Term | Benign | Malignant | Either | Notes |
|---|---|---|---|---|
| adenoma | X | | | |
| adenomocarcinoma | | X | | |
| anaplastic/anaplasia | | X | | poorly differentiated or undifferentiated cells that grow rapidly, malignant neoplasias often have anaplastic cells |
| angiolipoma | X | | | |
| angiomyolipoma | X | | | |
| angioma | X | | | |
| basal cell tumor | X | | | typically benign terminology, called basal cell carcinoma if malignant |
| blastoma | | X | | |
| carcinoma | | X | | suffix used to denote malignancy |
| chondroma | X | | | |
| chromatophoroma | | | X | |
| cystadenoma | X | | | |
| cytoma | | | X | just a suffix for tumor/cancer, depends on prefix and circumstances |
| dysgerminoma | | X | | |
| epulis | X | | | |
| fibroma | X | | | |
| fibrosarcoma | | X | | |
| granulosa cell tumor | | X | | |
| hemangioma | X | | | |
| hepatic osseous metaplasia | | | X | OM is bone-y growths that can occur in benign or malignant tumors |
| hepatoma | | X | | often used as shorthand to describe hepatic carcinoma |
| hyperplasia | X | | | |
| insulinoma | | | X | 90% benign, small chance of malignancy |
| leiomyoma | X | | | |
| leukemia | | X | | |
| leukosis | | X | | leukemia-like malignant viral disease seen especially in cattle and birds |
| lipoma | X | | | |
| lymphoma | | X | | benign lymphomas are rare and usually specified as benign in name |
| mast cell tumor | | | X | |
| melanoma | | X | | |
| melanophoroma | | | X | |
| meningioma | | | X | typically benign, small percent may become malignant |
| mesiothelioma | | | X | benign is less common but possible |
| myelolipoma | X | | | |
| myxosarcoma | | x | | malignant tumor of fibroblastic origin |
| neoplasia | | | X | |
| osteoma | X | | | |
| papilloma | X | | | |
| pheochromocytoma | X | | | small chance of spreading to other parts of the body |
| sarcoma | | X | | suffix used to denote malignancy |
| seminoma | | X | | |
| squamous cell carcinoma | | X | | |
| sertoli cell tumor | | | X | |
| teratoma | | | X | mature teratoma benign, immature may develop into malignant |

| | | | | |
|---|---|---|---|---|
| thymoma | | | X | high probability of becoming malignant |
| trichoepithelioma | X | | | |
| tumor | | | X | |
| xanthogranuloma | X | | | typically benign and self-limiting |
